# Brain monoaminergic activity during predator inspection in the Trinidadian guppy (*Poecilia reticulata*)

**DOI:** 10.1101/2021.11.25.469990

**Authors:** S. Dimitriadou, S. Winberg, P.-O. Thörnqvist, DP. Croft, SK. Darden

## Abstract

To understand the processes underpinning social decision-making, we need to determine how internal states respond to information gathered from the social environment. Brain monoamine neurotransmitters are key in the appraisal of the social environment and can reflect the internal state underlying behavioural responses to social stimuli. Here we determined the effects of conspecific partner cooperativeness during predator inspection on brain monoamine metabolic activity in Trinidadian guppies (*Poecilia reticulata*). We quantified the concentration of dopamine, serotonin and their metabolites across brain sections sampled immediately after ostensibly experiencing cooperation or defection from social partners whilst inspecting a predator model. Our results indicate dopaminergic and serotonergic activity differs with the cooperativeness experienced; these different neurotransmission profiles are likely to affect the expression and regulation of downstream behaviours that ultimately contribute to the patterning of cooperative interactions among individuals in the population.

List of abbreviations used

## 1 Introduction

Individuals continuously perform evaluation checks to stay current with factors in their environment important to their survival and reproduction. These checks include gathering information on intrinsic valence, novelty, and violation of expectations, to assist with evaluating the valence (positive/negative) and salience (high/low) of both stimuli and the resources (including coping mechanisms) available to the individual for dealing with them [1, 2]. This appraisal subsequently affects the animal’s internal state, or core-affect, as a function of their perception of the environment [3] and thereby drives future behaviour [4]. Quantifying the appraisal-to-behavioural response pathway can thus provide us with insight into decision-making processes and ultimately the rules and strategies that guide behavioural outputs [5]. Such an approach has great potential to understand the rules and strategies that govern decisions made in the context of real-world cooperative interactions, where appraisal of the social environment quite often is central, but such studies are rare.

The paradoxical existence of cooperation in populations where individuals cooperate with non-kin has received considerable scientific interest [6–8]. A number of rules and strategies guiding behavioural responses to the cooperative behaviour of others have been proposed in the theoretical literature to underpin its persistence, ranging from different forms of reciprocity of cooperative acts to rules of association and punishment of defectors [8–11]. However, we currently have very little understanding of the individual decision-making processes that lead individuals to modify their behaviour according to information gleaned regarding the cooperativeness of others, such as the decision to reciprocate or not the behaviour of partners [e.g. 12,13] or to continue or not any particular social affiliation [e.g. 14,15]. Unravelling the mechanistic underpinnings of the appraisal of the social environment will provide insight into cooperative decision-making processes and ultimately the rules and strategies that maintain cooperation among unrelated individuals.

Monoamine neurotransmitters, such as dopamine (DA), norepinephrine (NE) and serotonin (5-hydroxytryptamine, 5-HT), have been shown to modulate numerous behaviours and physiological functions, including attention [16, 17], reward and risk assessment [18–20], stress responses [21, 22] mood, emotion and fear [23, 24]. In fish and other vertebrates monoaminergic neurotransmission has been directly implicated in social interaction and social stress [25–32] and in decision-making processes [33–35]. This may be due to the involvement of monoamine neurotransmitters in processes underlying stimulus appraisal and therefore in core affect or emotion-like states, including reward and prediction error, motivation, arousal, brain affect, emotional bias, and emotional memory [19,20,23,36–39].

Given these documented roles of monoaminergic neurotransmission, studies quantifying their involvement in social decision-making processes associated with cooperative interactions are well-warranted. To date, however, past research has focused on the effect of monoaminergic neurotransmission on the *expression* of cooperative behaviour and has been largely limited to investigations of heterospecific cooperation [35,40,41]. We currently have a gap in knowledge regarding the *response* of these systems to variation in conspecific cooperation. Consequently, we have only a limited understanding of the psychological processes triggered by these experiences and their downstream behavioural effects.

In the current study, we use the Trinidadian guppy (*Poecilia reticulata*) as a model system to quantify how the cooperativeness of social partners affects brain neurotransmission. Guppies cooperate during predator inspection, a behaviour in which a small group of fish leave the relative safety of the shoal or other refuge to approach a potential predator and assess the level of threat posed; they then return to the shoal and transmit this information [42–44]. Predator inspection is considered a model for the study of cooperation [45], as all shoal members benefit from the information gathered, irrespective of whether they inspected or not. Previous work suggests that brain nonapeptide production [46] and downstream behaviours [15, 47] are measurably affected by the previous experience of the cooperative acts of others. We experimentally manipulated the ostensible experience of cooperation or defection from the social environment, and then measured the brain levels of DA, 5-HT, and their metabolites. We found that monoaminergic activity differed between brain sections with treatment, indicative of changes in internal state that may mediate the downstream effects of experiencing cooperation or defection. The transmission patterns we explore represent a first and key step in understanding the psychological mechanisms underpinning conditional cooperation in non-human animals and more generally, provide insight into the mechanisms underlying cooperation among non-kin.

## 2 Materials and methods

### 2.1 Study subjects

One hundred and twenty juvenile (sexually immature) Trinidadian guppies, descendants of wild-caught fish from a high predation site of the Aripo River on the island of Trinidad (10°39′27N, 61°13′34W), were collected from mixed-generation pools in the University of Exeter, Department of Psychology fish laboratory facilities for rearing in a standardized environment [tank dimensions: 80x30x40 cm; 12h light: 12h dark cycle]. The fish were fed with commercial flake and live food (*Artemia* sp.) twice a day and were kept in a constant room temperature of 25°C. Upon reaching sexual maturity 56 females were tested. Stimulus fish originated from the same population and were kept in the same conditions as focal fish.

All experiments were undertaken under a U.K. Home Office project licence (P5786D4EA) and a personal licence (I002BDF3F), and were in accordance with the UK Animals (Scientific Procedures) Act, 1986 and the ARRIVE guidelines.

### 2.2 The cooperation paradigm

Experience of the behaviour of social partners in a cooperative context was manipulated in female Trinidadian guppies (N=56; 8 fish were excluded from the study because of issues with the scoring of their behaviour through video recording; these were distributed across experimental conditions) using a predator inspection paradigm similar to those commonly used for small freshwater fish [48–51]. For our experimental condition this entailed presenting free-swimming focal individuals with a predatory stimulus (a realistic model of a pike cichlid, *Crenincichla alta*) at the end of an inspection lane (Fig. S1) and allowing them to inspect. During control trials, focal fish were presented with a plastic aquarium plant (see below for rationale for the two conditions). Fish were tested singly, but were provided with a same-sex, size-matched shoal of 4 conspecifics, who were constrained behind a transparent, perforated Perspex barrier for visual and olfactory contact with the focal individual, but could not perform an inspection themselves. To simulate cooperation (joining by social partners) one side of each inspection lane was lined with a mirror, allowing the focal individual to inspect with their mirror image [51]. Defection (not joining by social partners) was simulated with an opaque surface lining the inspection lane. The simulation of cooperation and defection through the use of mirrors in this paradigm has been widely used in this species [45,49,50,52] and it has been recently demonstrated that the cooperative behaviour of guppies in this context is highly correlated with individual cooperativeness measured in predator inspection trials with live partners [51]. In addition, this experimental approach has been recently used to look at the effects of the manipulation of neurotransmission on cooperative behaviour [53].

Each focal individual was assigned to either an experimental (predator model) or a control (plastic plant) condition. The experimental condition ostensibly manipulated whether focals experienced their social partners as cooperating or defecting during predator inspection, while the control condition replicated all aspects of the experimental environment that may have affected neurotransmission patterns outside of the cooperative context (i.e. inspection of a predator), including effects of having a mirrored or non-mirrored lane (e.g., perception of lane size). On the latter point, we refer to the control treatment where a social partner is simulated through the use of a mirror as ‘cooperation’, even though the inspection of the control stimulus is not necessarily a cooperative act, as there is no threat present. Instead of using live predators as inspection stimuli, we used realistic predator models of *Crenicichla frenata* (total length: 12cm), a common predator of adult guppies in the wild [54, 55]. Predator models are widely used for predator inspection studies in the literature [44,56,57] because they elicit an anti-predator response and offer standardised predator behaviour, thus eradicating confounds introduced by variation in the behaviour of live predator stimuli.

A stimulus shoal consisting of 4 size-matched female conspecifics not previously encountered by focal fish was introduced in each stimulus shoal compartment. After a 20-minute time period, which allowed for the accumulation of olfactory cues as well as the acclimation of the stimulus shoal, a focal fish was introduced in the testing compartment and was left for 10 minutes to acclimatise. The focal fish had visual and olfactory access to the stimulus shoal throughout this period. At the end of the 10 minutes, when the focal fish entered the refuge area of its own accord, two visual barriers were lifted, uncovering the mirror (or an opaque surface for the defection groups) and the inspection stimulus. This signified the start of the up to 5-minute long experimental trial, during which the focal individual was free to inspect the inspection stimulus. The trial ended after one inspection (which was defined as the fish approaching the inspection stimulus compartment within a distance less than 22 cm and then returning to the refuge area), or after a 5-minute period if no inspection occurred (data from individuals that did not perform an inspection were excluded from the analysis). At the end of the trial, the focal fish was removed from the tank, and rapidly euthanised using ice slurry (maximum temperature of 4°C). Their brain was subsequently removed and dissected into three macro-areas: fore-section (including the telencephalon and the preoptic area, excluding the olfactory bulbs and the hypothalamus) (N=45), mid-section (including the optic tectum, diencephalon, and the hypothalamus) (N=42) and hind-section (cerebellum and medulla oblongata) (N=41) (see Supplementary material, Fig. S2). These macro-areas were used rather than traditionally defined regions because we were not able to reliably section the hypothalamus in these traditional regions in our samples. Each brain sample was stored in a 1.5 ml Eppendorf tube and instantly frozen at -80°C within 3 minutes of euthanasia.

Trials were video recorded, and videos were analysed using the Noldus Observer XT software (Wageningen, The Netherlands). The behavioural measures recorded were the distance of closest approach to the stimulus compartment (measured from the stimulus compartment, i.e. 0 cm correspond to the closest inspection) and the duration of an inspection (i.e. the time a focal individual spent inspecting the stimulus) (Supplementary Fig. S2, S3). Individuals who did not leave the refuge area were excluded from subsequent analysis (N=2); the same was true for individuals that approached the stimulus compartment at a distance smaller than 22 cm but did not perform an inspection and were exhibiting escape behaviour (fast swimming alongside the tank wall) (N=1).

### 2.3 Analysis of brain monoamines and protein content

Brain levels of 5-HT and its metabolite 5-hydroxyindoleacetic acid (5-HIAA), DA and DA metabolites 3,4-dihydroxyphenylacetic acid (DOPAC) and homovanillic acid (HVA), as well as NE were analysed using high performance liquid chromatography with electrochemical detection (HPLC-EC), using the same protocol as Thörnqvist, Höglund, and Winberg [58]. In brief, the frozen sectioned brain samples were homogenised in 4% (w/v) ice-cold perchloric acid, containing 10ng/ml 3,4-dihydroxybenzylamine (DHBA, internal standard), with the use of a Sonifier cell distributor B-30 (Branson Ultrasonic, Danbury, CT, USA) and were subsequently centrifuged at 21,000g for 10 minutes at 4°C. The supernatant was used for HPLC-EC in order to analyse the monoamine content of the samples, while the pellet was stored at -20°C for analysis of the protein content. The HPLC-EC system consisted of a solvent delivery system model 582 (ESA, Bedford, MA, USA), an autoinjector Midas type 830 (Spark Holland, Emmen, The Netherlands), a reverse phase column (Reprosil-Pur C18-AQ 3 μm, 100x4 mm column, Dr Maisch HPLC GmbH, Ammerburch-Entrigen, Germany) kept at 40°C and an ESA 5200 Coulochem II EC detector (ESA, Bedford, MA, USA) with two electrodes at reducing and oxidising potentials of -40 and +320 mV. In order to oxidise any contaminants, a guarding electrode with a potential of +450 mV was employed before the analytical electrodes. The mobile phase consisted of 75 mmol/l sodium phosphate, 1.4 mmol/l sodium ocyl sulphate and 10 μmol/l Ethylenediaminetetraacetic acid (EDTA) in deionised water containing 7% acetonitrile (pH 3.1, using phosphoric acid). The monoamine content of each sample was quantified by comparison with standard solutions of known concentrations. Correction for recovery was made with the use of DHBA as the internal standard, with the use of the HPLC software Clarity™ (DataEpex Ltd, Prague, Czech Republic). For normalisation of brain monoamine levels, the concentration of total protein in the brain sample was used.

To assess protein content, the pellets of the centrifuged, homogenised brain sections were diluted in 100 μl of Tris(hydroxymethyl)aminomethane (Tris) buffer, using a Sonifier cell distributor B-30 (Branson Ultrasonic) to ensure full dilution of the pellet. A QuBit 2.0 Fluorometer (InVitrogen, Carlsbad, CA) was used to analyse the protein concentration, by measuring absorbance at 280nm. The concentration of monoamines and their metabolites was expressed as ng per mg of protein [59]. The ratio of the concentration of the metabolite to that of the parent monoamine in the tissue was used for all subsequent analysis, as it is found to be a good indicator of neural activity (higher metabolite-to-monoamine ratios show increased release and turnover rates of the corresponding neurotransmitters) [59, 60]. The turnover ratio of norepinephrine could not be calculated because of technical difficulties at detecting its metabolites with the methodology used. Samples where the quantity of neurotransmitters and/or metabolites could not be confidently calculated were excluded from the analysis (fore-section: N= 3, mid-section: N= 4; hind-section: N=2).

The analysis of brain monoamines and protein content took place at the Department of Neuroscience of the University of Uppsala (Biomedical Center).

### 2.4 Statistical analysis

Whole brain monoamine turnover rates (concentration of metabolite/concentration of parent monoamine) were analysed by fitting linear models for DOPAC/DA and 5-HIAA/5-HT turnover ratios after logarithmic transformation. HVA/DA turnover rates were analysed using beta regression in the ’betareg’ v3.0-5 R package [61]. Beta regression allows statistical modelling of continuous, restricted to the unit interval (0,1), non-transformed data [62]. Monoamine turnover rates were also analysed separately for every brain section (linear models for the logarithm DOPAC/DA and 5-HIAA/5-HT turnover rates; beta regression for HVA/DA rates). All statistical analyses were carried out in R v3.2 [63]. To control for effects of the inspection behaviour of the focal individual on monoamine activity, the distance of closest approach to the stimulus and the time spent inspecting the stimulus were included in the model. In all cases the full model included Standard body length + Distance of closest approach during inspection + Duration of stimulus inspection + Social Environment (Cooperation/Defection) + Inspection stimulus (Control/Predator) + Social Environment* Inspection stimulus.

## 3 Results

### 3.1 Whole brain monoaminergic activity is affected by experiencing cooperation from the social environment

Across the whole of the brain, log-transformed DOPAC/DA ratios tended to be affected by the interaction of the social environment (ostensible cooperation versus defection) and the experimental condition (predator versus control) [two way interaction: F(1,38)=3.699, p=0.062] (Fig. 1A); this trend, however, did not reach statistical significance (Table 1). A similar trend was observed for log transformed 5-HIAA/5-HT ratios (Fig. 1B) (Table 1). Whole brain 5-HIAA/5-HT ratios (after log transformation) were found to depend on the social experience during the behavioural trial (Fig. 1B), with fish experiencing cooperation showing lower 5-HIAA/5-HT ratios than those experiencing defection [F(1,37]=6.875, p=0.013], irrespective of whether this was in the presence of a predator or a plant stimulus. HVA/DA ratios were found to be independent of these factors (data not shown) (Table 1). The focal individual’s standard body length, distance of closest approach to the predator compartment and duration of the behavioural trial were found to have no effect on whole brain monoamine turnover rates (Table 1).

**Fig. 1.**
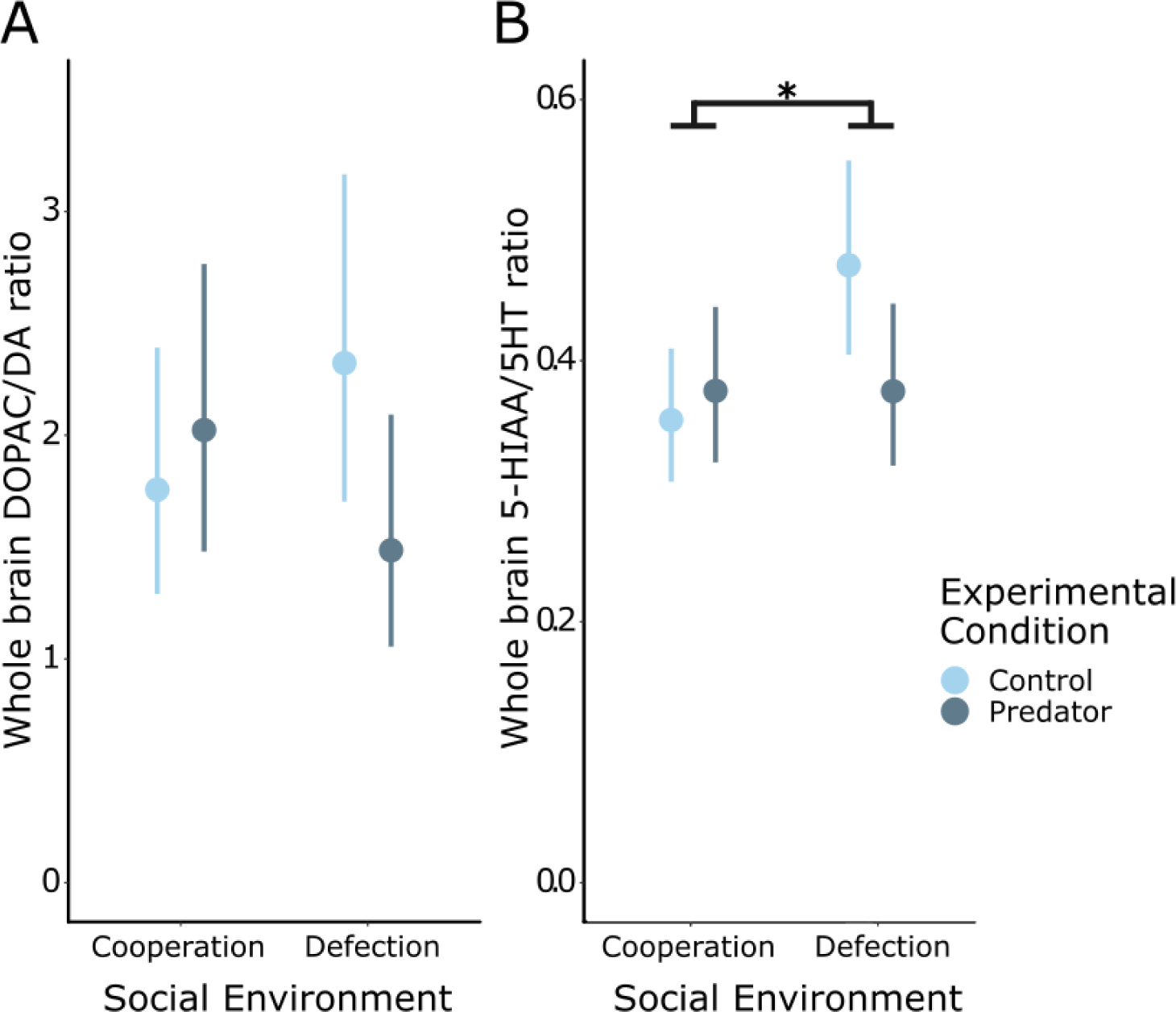
Effects of cooperation or defection during predator inspection in log-transformed whole brain monoamine metabolism (A: DOPAC/DA; B: 5-HIAA/5-HT). A. DOPAC/DA ratios were not affected by the interaction of the social environment and the experimental condition. B. Log-transformed 5-HIAA/5-HT ratios were found to be affected by the social environment, with fish experiencing cooperation showing lower 5-HIAA/5-HT ratios than those experiencing defection. Back-transformed estimated marginal means and 95% confidence intervals (Cooperation-Control: N= 10; Cooperation-Predator: N= 9; Defection-Control: N= 11; Defection-Predator: N= 8). ** p< 0.01

**Table 1.**
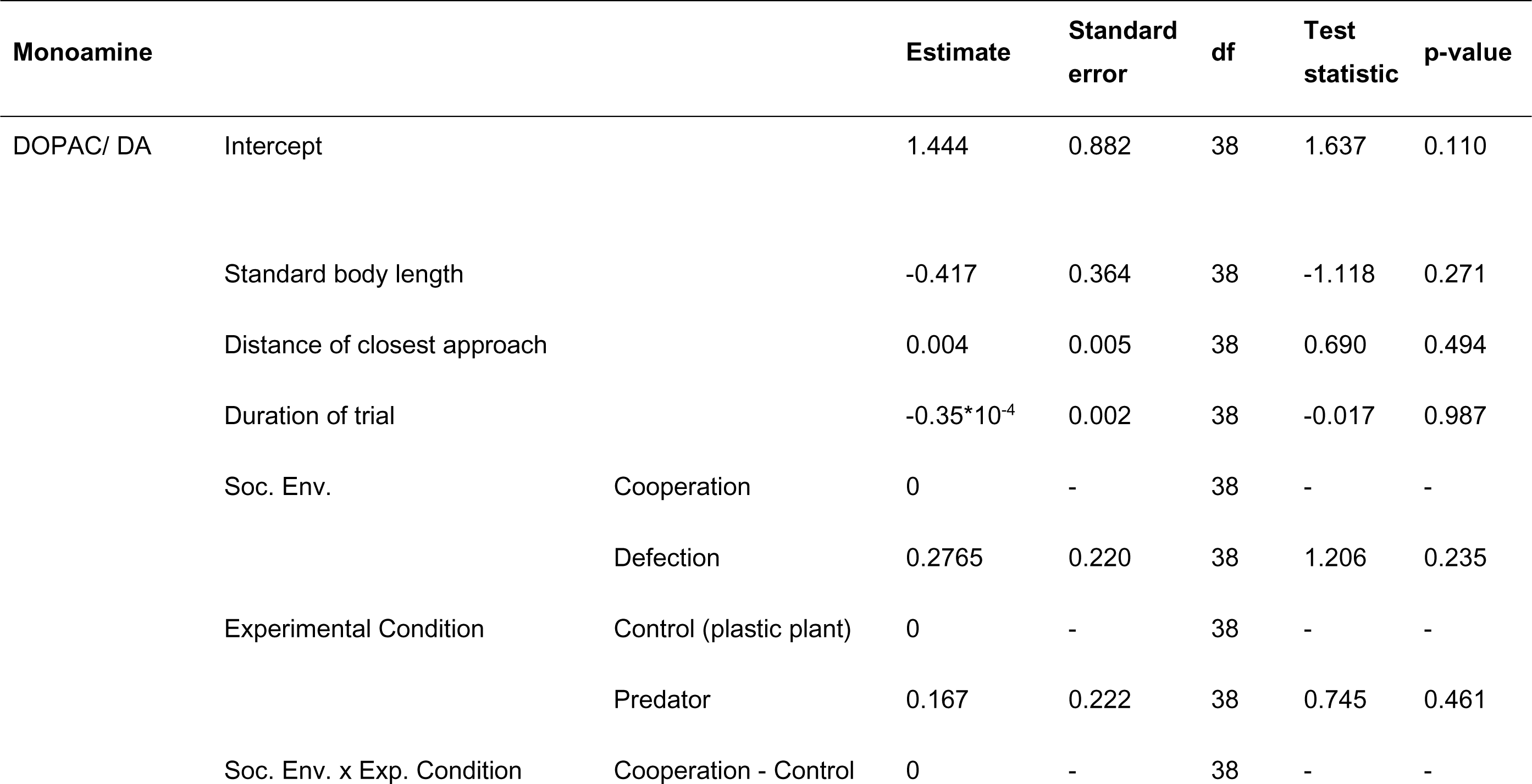
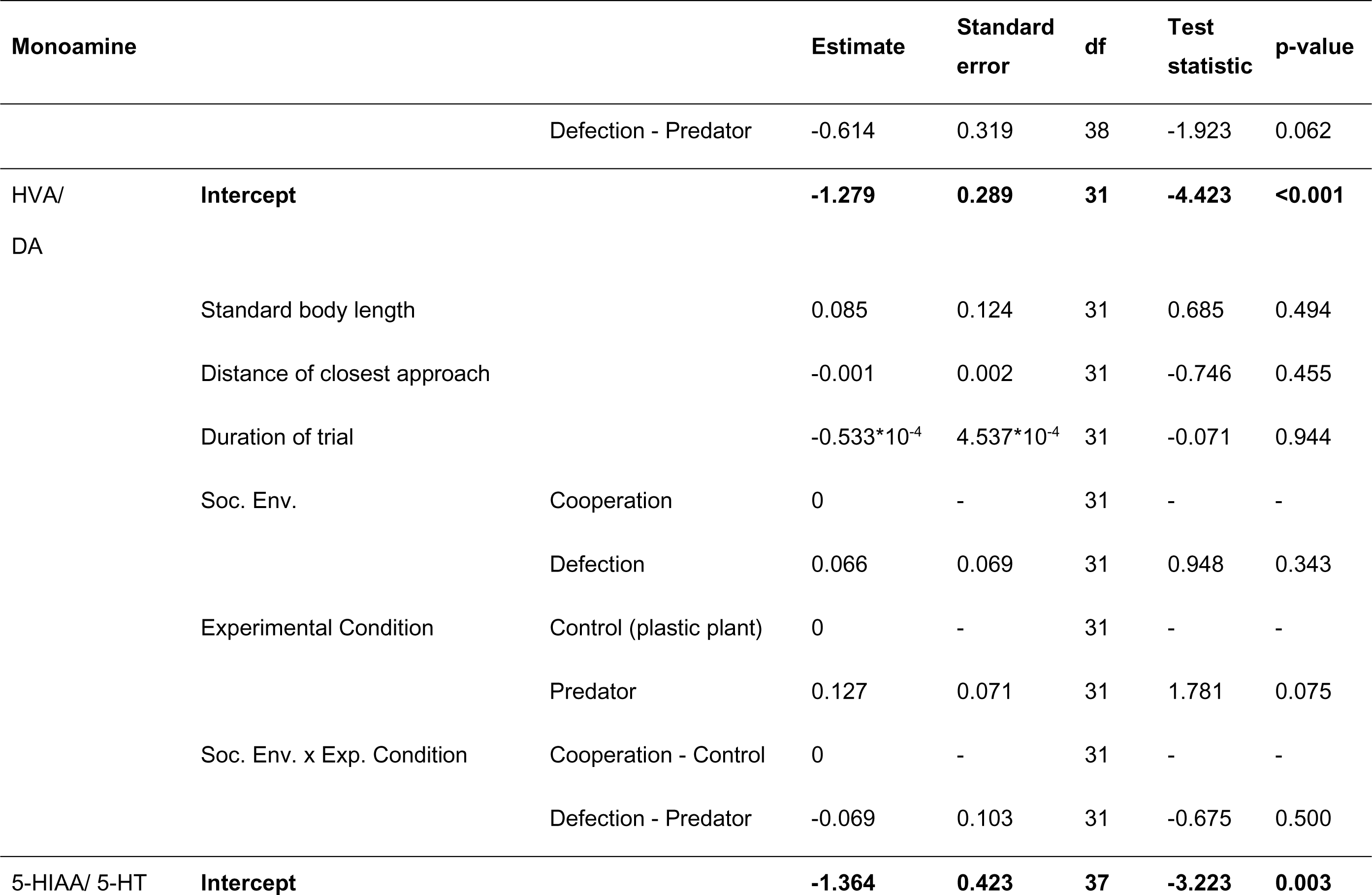
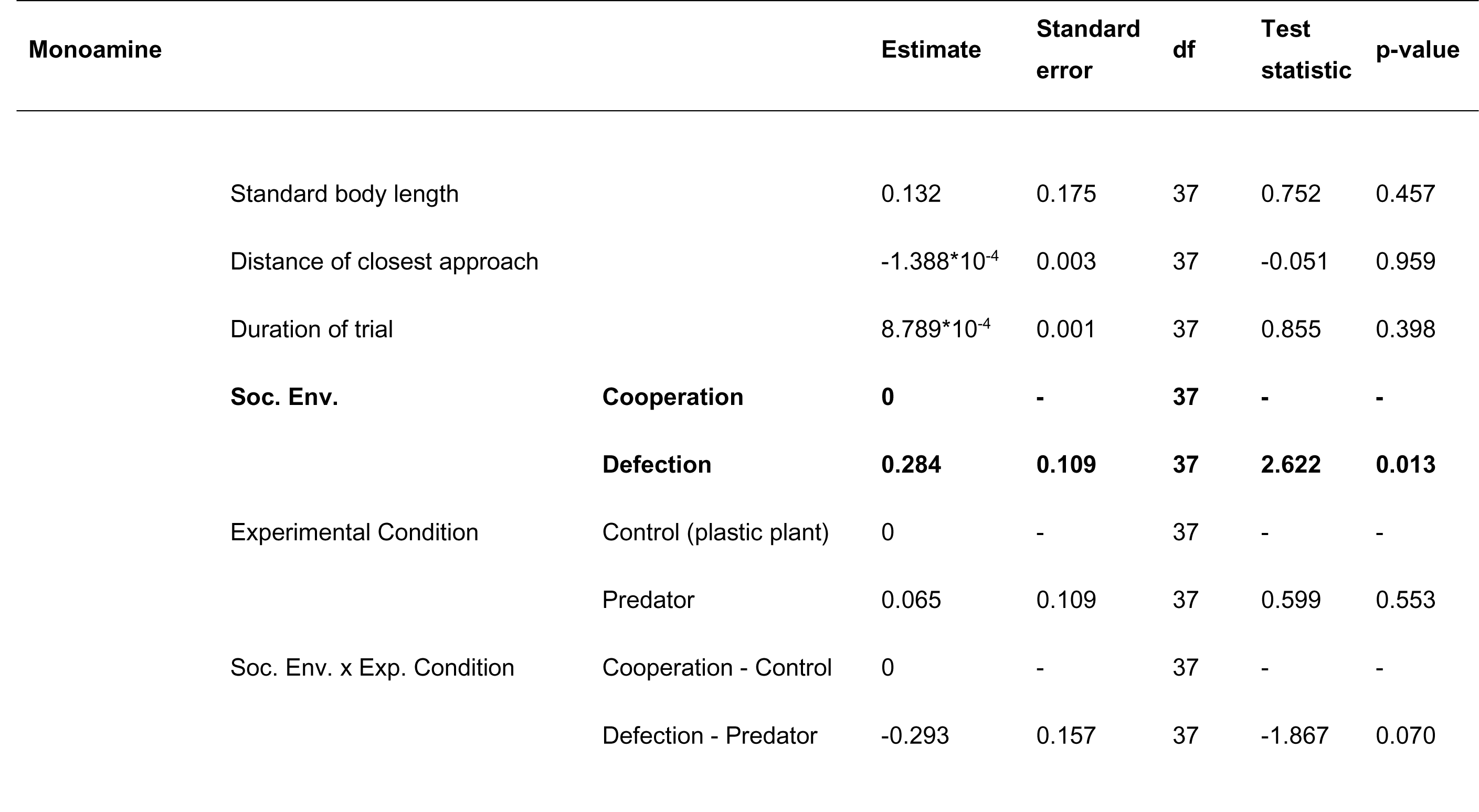
Marginal effects of standard body length, distance of closest approach to the predator compartment, duration of trial, social environment and experimental condition on whole brain neurotransmitter turnover rates (DOPAC/DA and 5-HIAA/5-HT after log transformation; HVA on non-transformed data). Statistically significant factors are shown in bold.

### 3.2 Experiencing cooperation affects neurotransmission differentially across the brain

To better understand the response of different brain areas to ostensibly experiencing cooperation or defection from social partners, neurotransmitter turnover rates were analysed separately for each of the three brain sections. Hind-section DOPAC/DA ratios were affected by the interaction of social experience (ostensible cooperation versus defection) and experimental condition (predator versus control) [two-way interaction: F(1,40)=5.360, p=0.026]. Post hoc analysis showed that in the absence of a cooperating social partner, inspecting a predator led to lower DOPAC/DA ratios than the corresponding control condition (Fig. 2C) (Table 3). We found no significant effect of experimental condition and/or ostensible experience of cooperation on the logarithm of the DOPAC/DA ratio in the fore-section and mid-section of fish (Fig. 2A, 2B) (Table 2). Focal individual standard body length, distance of closest approach to the predator and trial duration had no effect on the DOPAC/DA ratio in any of the brain sections (Table 2).

**Fig. 2.**
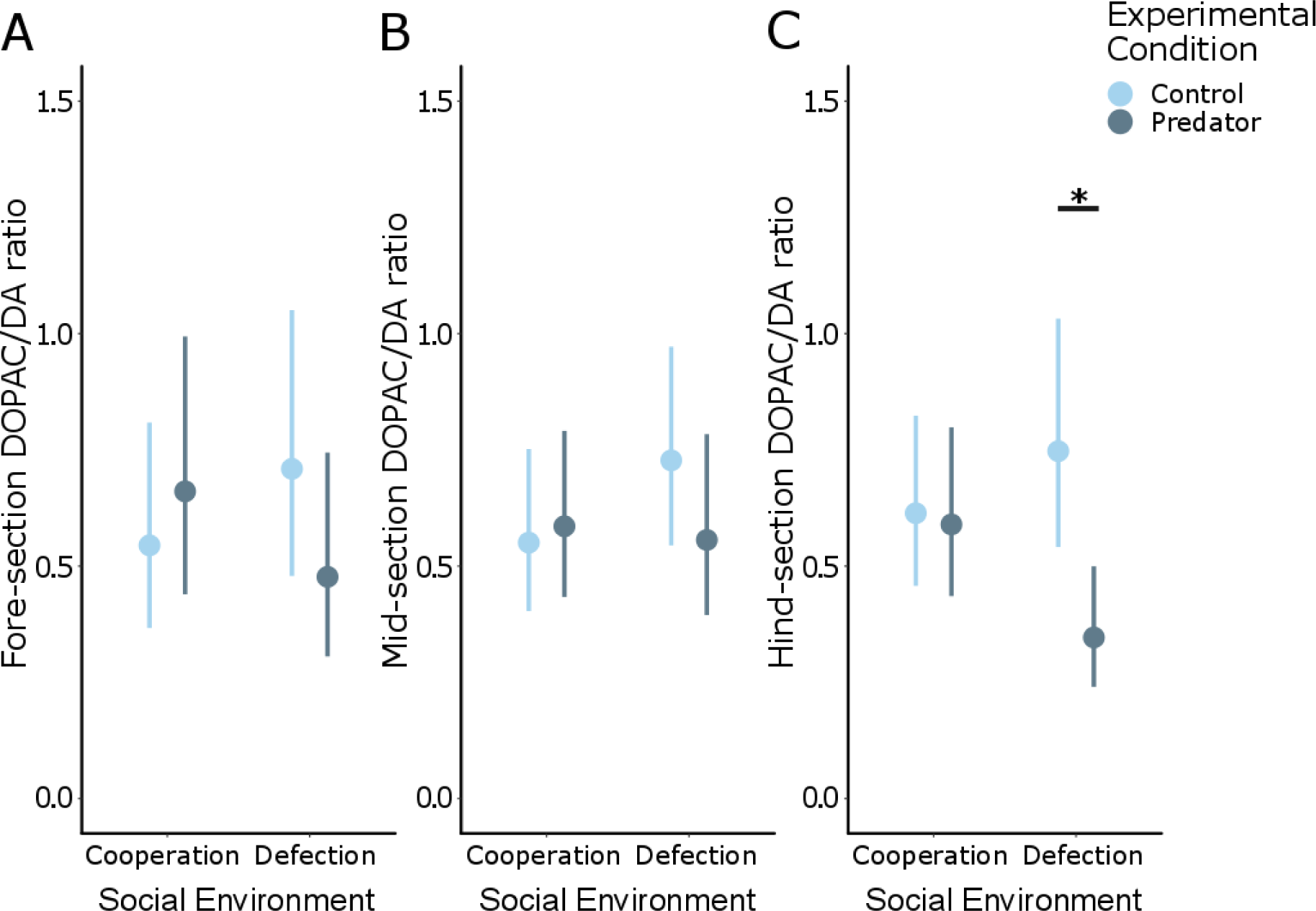
Effects of experiencing cooperation or defection during predator (dark grey) exposure or exposure to a plastic plant (light blue) on the log transformed DOPAC/DA ratios in the fore-section (A), mid-section (B), and hind-section (C) of Trinidadian guppies. A. We found no significant effect of experimental condition and/or ostensible experience of cooperation on the logarithm of the DOPAC/DA ratio in the fore-section of fish (Cooperation-Control: N= 14; Cooperation-Predator: N= 12; Defection-Control: N= 13; Defection-Predator: N= 10). B. Mid-section DOPAC/DA ratios were not affected by the cooperative behaviour of the social environment, or by the experimental condition (Cooperation-Control: N= 12; Cooperation-Predator: N= 12; Defection-Control: N= 14; Defection-Predator: N= 10). C. When focal individuals experienced defection from their social environment, predator inspection led to lower hind-section DOPAC/DA ratios than the control (plant) condition (Cooperation-Control: N= 13; Cooperation-Predator: N= 12; Defection-Control: N= 14; Defection-Predator: N= 11). Back-transformed estimated marginal means and 95% confidence intervals. * p< 0.05

**Table 2.**
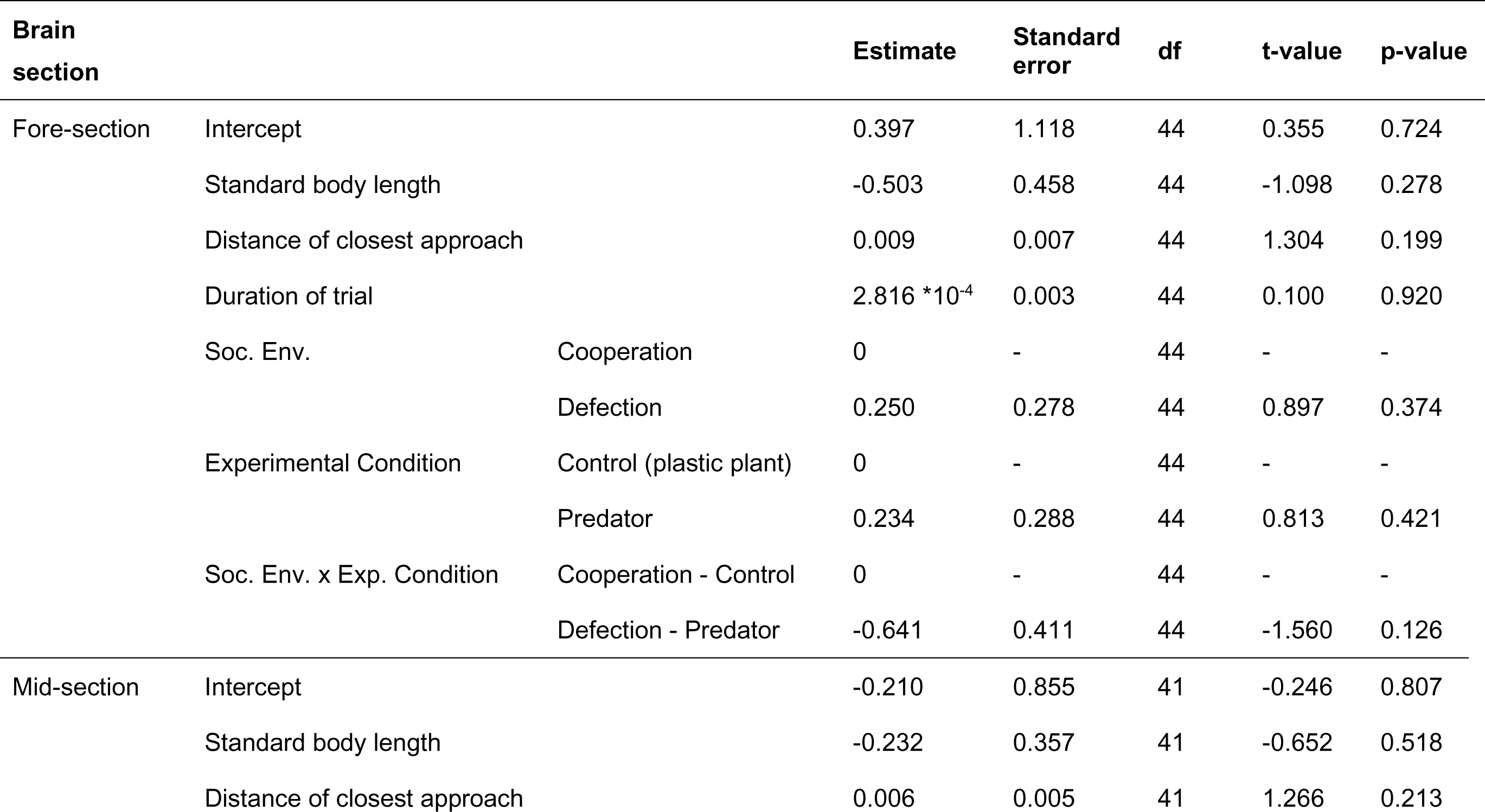
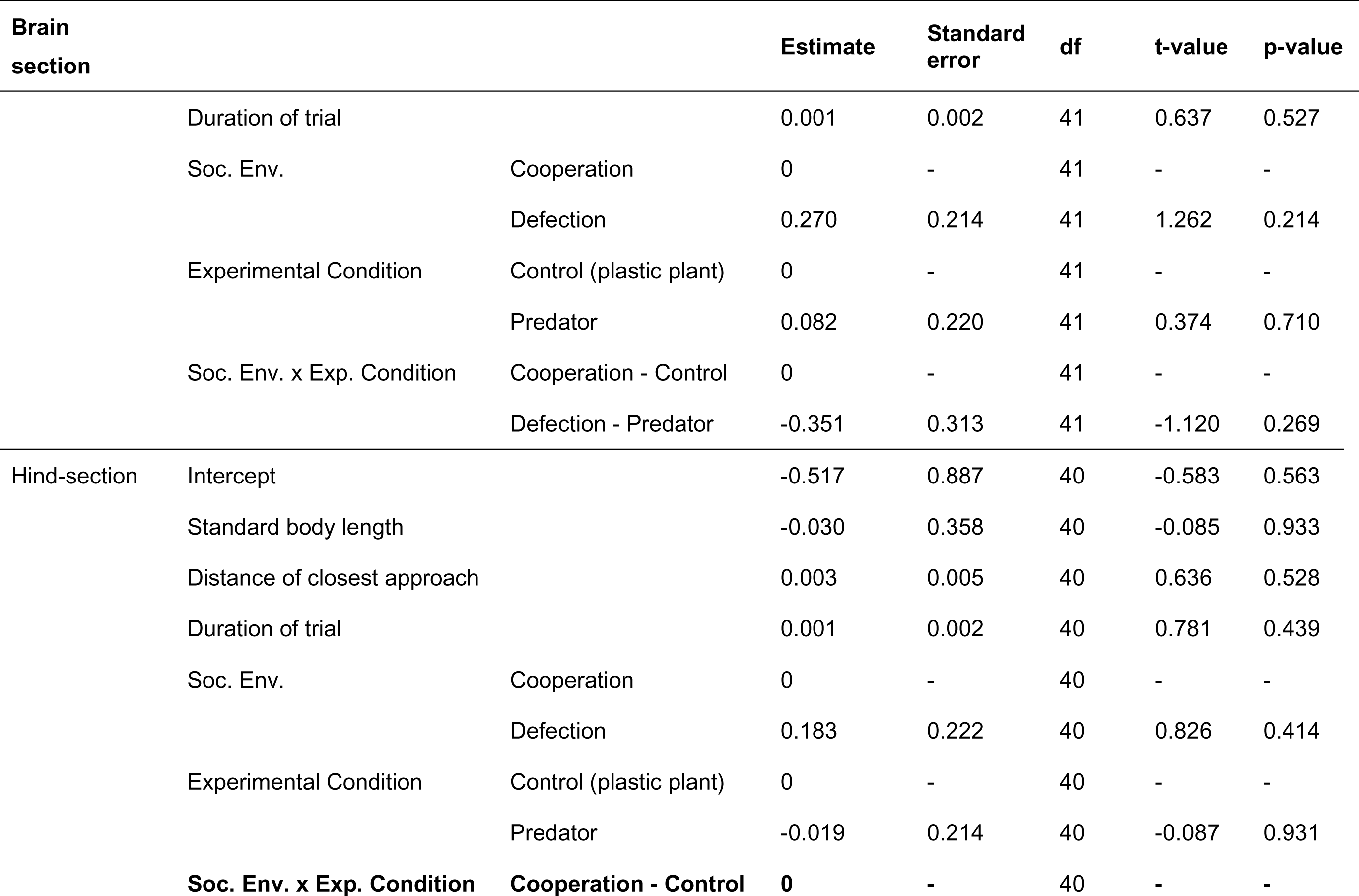
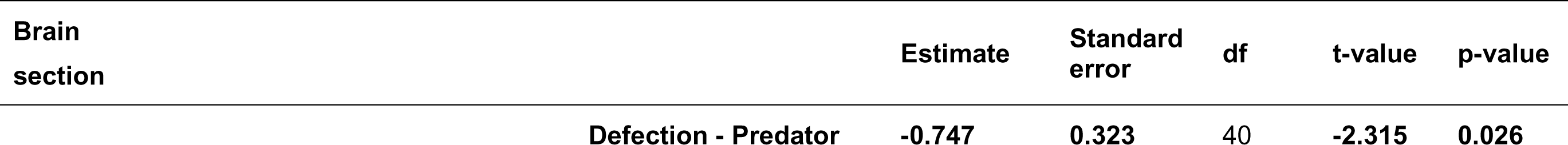
Marginal effects of standard body length, distance of closest approach to the predator compartment, duration of trial, social environment, and experimental condition on the log-transformed DOPAC/DA ratio in the fore*-section*, mid*-section* and hind*-section*. Statistically significant factors are shown in bold.

**Table 3.** Post hoc analysis for the ‘Social Environment x *Experimental* Condition’ interaction on the logarithm of the hind-section DOPAC/DA ratio. Pairwise least squares means comparisons. Statistically significant contrasts are shown in bold.

| Contrast | Estimate | Standard error | df | t-value | p-value |
| --- | --- | --- | --- | --- | --- |
| Coop. Control – Def. Control | -0.183 | 0.222 | 40 | -0.826 | 0.842 |
| Coop. Control – Coop. Predator | 0.019 | 0.214 | 40 | 0.087 | 0.999 |
| Coop. Control – Def. Predator | 0.582 | 0.237 | 40 | 2.461 | 0.082 |
| Def. Control – Coop. Predator | 0.202 | 0.227 | 40 | 0.889 | 0.810 |
| **Def. Control – Def. Predator** | **0.766** | **0.241** | 40 | **3.182** | **0.014** |
| Coop. Predator – Def. Predator | 0.564 | 0.242 | 40 | 2.327 | 0.108 |

Fish that had ostensibly experienced cooperation during a trial showed higher hind-section HVA/DA ratios than those in the defection treatment, irrespective of experimental condition (predator versus control) [F(1,37)=4.779, p=0.029] (Fig. 3C). We found a non-significant trend for distance of closest approach to the predator compartment to affect HVA/DA ratio in the hind-section [χ^2^(1, 37)=2.925, p=0.087), with fish approaching the predator compartment more closely tending towards lower HVA/DA ratios. DA to HVA turnover rates were found to be independent of experimental condition and the ostensible experience of cooperation or defection in the fore-section (Fig. 3A) and mid-section (Fig. 3B) of the tested fish. Focal individual standard body length did not affect HVA/DA ratios in any of the brain sections (Table 4).

**Fig. 3.**
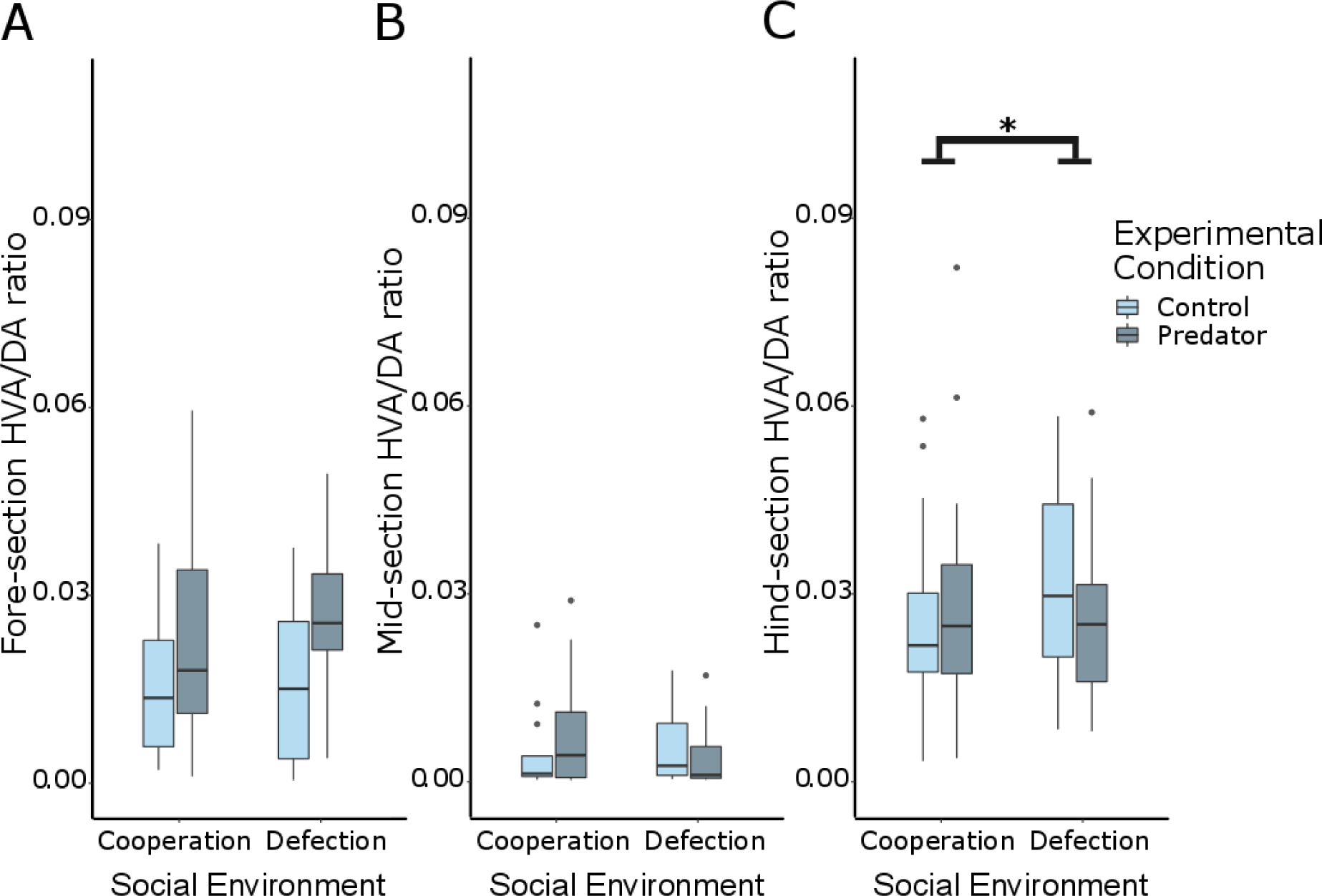
HVA/DA ratios in the fore-section (A), mid-section (B), and hind-section (C). A. Fore-section HVA/DA ratios were not affected by the cooperative behaviour of the social partners, or by the experimental condition (Cooperation-Control: N= 14; Cooperation-Predator: N= 12; Defection-Control: N= 13; Defection-Predator: N= 10). B. DA to HVA turnover rates were found to be independent of experimental condition and the ostensible experience of cooperation or defection in the mid-section (Cooperation-Control: N= 12; Cooperation-Predator: N= 12; Defection-Control: N= 14; Defection-Predator: N= 10). C. Experiencing cooperation by the social environment led to higher hind-section HVA/DA ratios than experiencing defection (Cooperation-Control: N= 13; Cooperation-Predator: N= 12; Defection-Control: N= 14; Defection-Predator: N= 11). Boxes represent the interquartile range (25th and 75th quartiles), and the horizontal lines represent the medians. The whiskers extend to the largest value (upper whisker) and lowest (lower whisker) value no further than 1.5 times the interquartile range. The dots represent outlying values. * p< 0.05

**Table 4.**
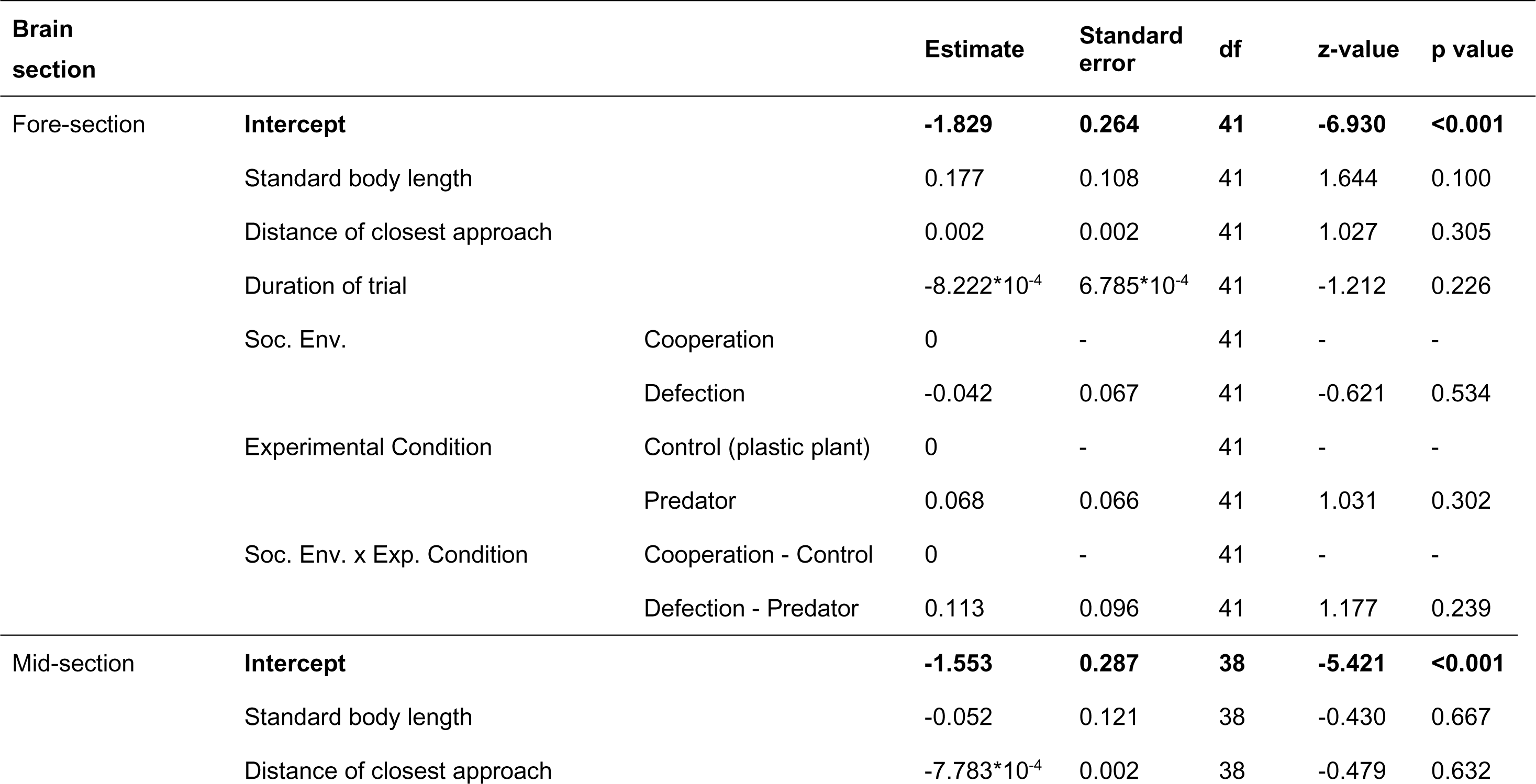
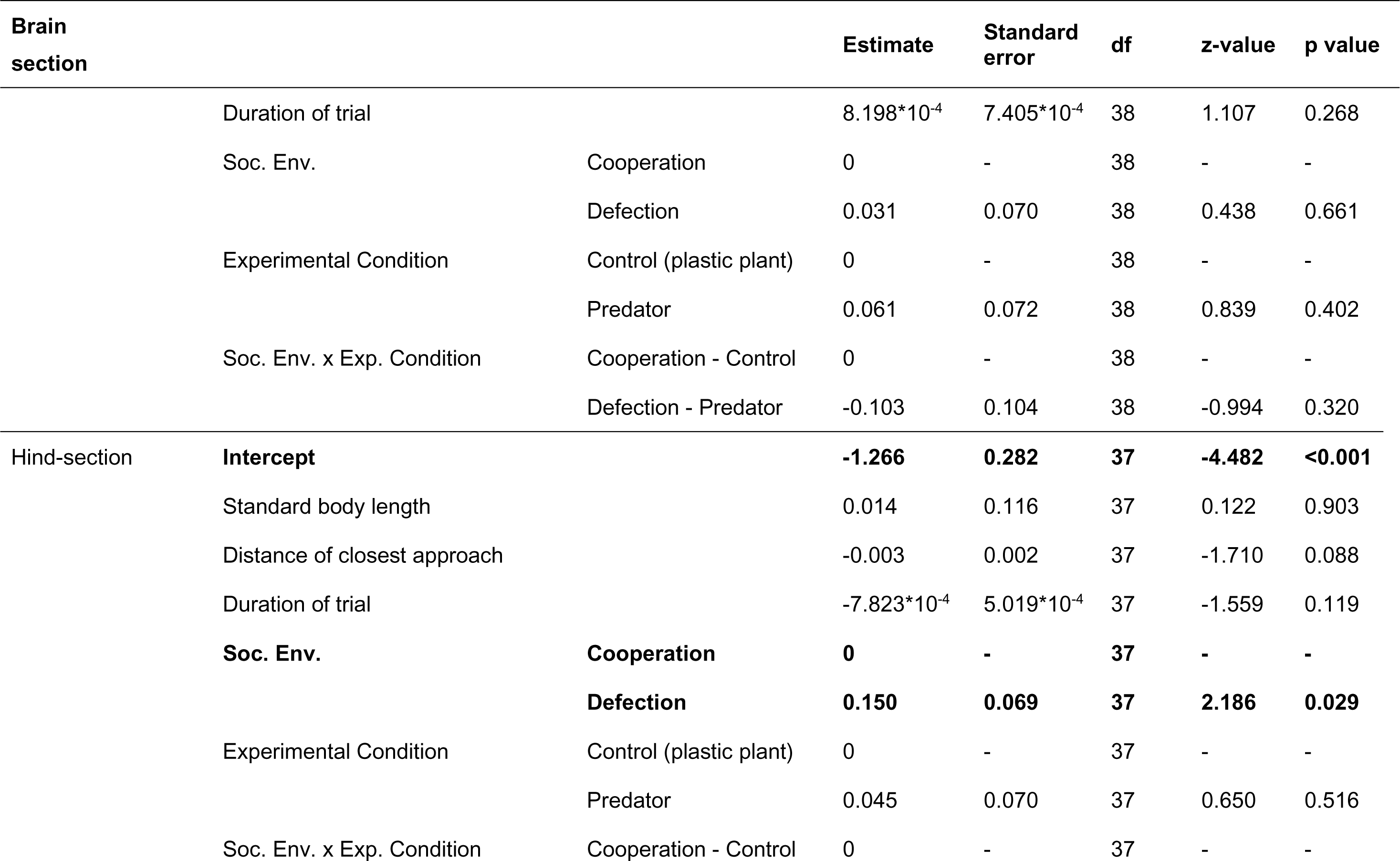
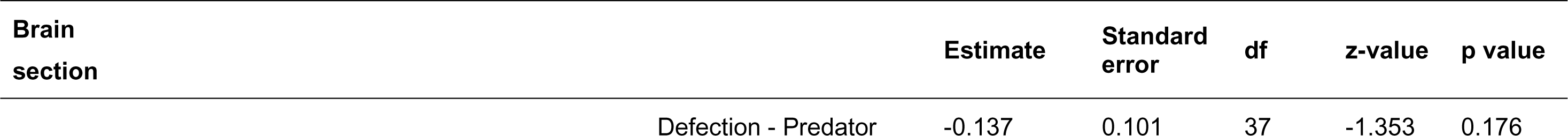
Marginal effects of distance of standard body length, closest approach to the predator compartment, trial duration, social environment and experimental condition on the HVA/DA ratio in the fore-section, mid-section and hind-section. Statistically significant factors are shown in bold.

Fore-section log-transformed 5-HIAA/5-HT ratios were affected by the interaction between the social environment during predator inspection (ostensible cooperation versus defection) and experimental condition (predator versus control) [two-way interaction: F(1,42)=4.390, p=0.042] (Fig. 4A) (Table 5). Post hoc analysis did not show statistically significant differences between pairs (Table 6); however, visual interpretation of the effect would suggest that this is driven by a crossover effect between the cooperative behaviour of the social partners (cooperation/defection) and the experimental condition (control/predator) (Fig. 4A) and would likely have been picked up with larger statistical power (current statistical power ∼61% at the 0.05 significance level). Ostensible social experience had a significant effect on log transformed 5-HIAA/5-HT ratios in the hind-section, irrespective of experimental condition (predator versus control) [F(1,40)=7.085, p=0.011], with fish ostensibly experiencing defection showing higher 5-HIAA/5-HT ratios than those ostensibly experiencing cooperation (Fig. 4C) (Table 5). Mid-section 5-HIAA/5-HT was independent of these factors (Fig. 4B). Focal individual standard body length, distance of closest approach to the predator compartment and duration of inspection had no effect on 5-HIAA/5-HT ratios in any of the brain sections studied (Table 5).

**Fig. 4.**
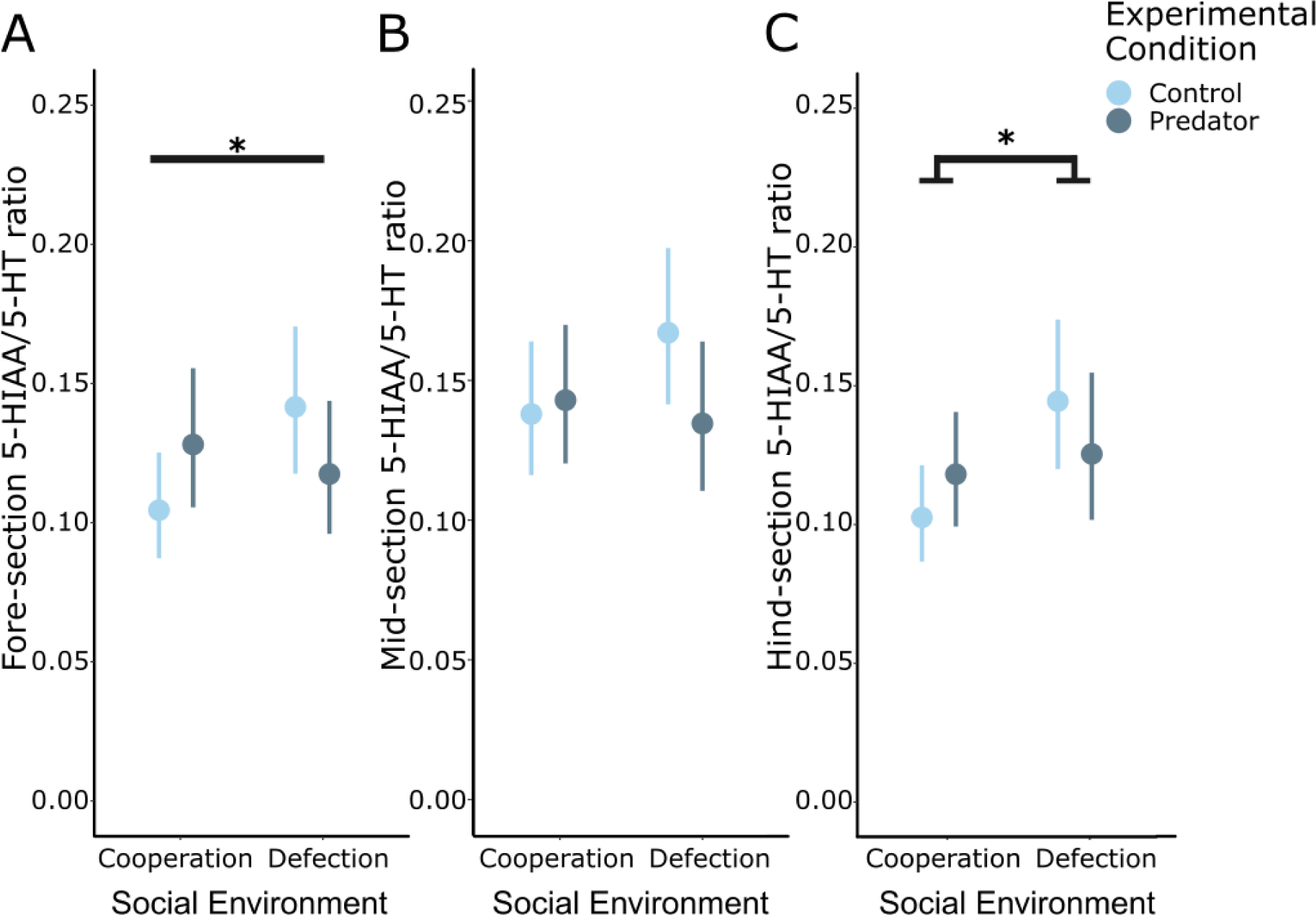
Serotonin metabolism rates (5-HIAA/5-HT) in the fore-section (A), mid-section (B), and hind-section (C) of guppies. A. Log-transformed fore-section 5-HIAA/5-HT ratios were affected by the interaction between the social environment and the experimental condition (predator versus control condition). Post hoc analysis did not show statistically significant differences between conditions (Cooperation-Control: N= 14; Cooperation-Predator: N= 12; Defection-Control: N= 13; Defection-Predator: N= 10). B. The cooperative behaviour of the social environment and the experimental condition had no effect on mid-section 5-HIAA/5-HT ratios (Cooperation-Control: N= 12; Cooperation-Predator: N= 12; Defection-Control: N= 14; Defection-Predator: N= 10). C. Experiencing defection led to higher hind-section 5-HIAA/5-HT ratios compared to experiencing cooperation across experimental conditions (Cooperation-Control: N= 13; Cooperation-Predator: N= 12; Defection-Control: N= 14; Defection-Predator: N= 11). Back-transformed estimated marginal means and 95% confidence intervals. * p< 0.05; ** p< 0.01

**Table 5.** Marginal effects of standard body length, distance of closest approach to the predator compartment, duration of behavioural trial, social environment and experimental condition on the log-transformed serotonin turnover rate (5-HIAA/5-HT) *in* the fore-section, mid-section and hind-section. Statistically significant factors are shown in bold.

| Brain section |  | Estimate | Standard error | df | z-value | p value |
| --- | --- | --- | --- | --- | --- | --- |
| Fore-section | **Intercept** | **-2.448** | **0.514** | **42** | **-4.763** | **<0.001** |
|  | Standard body length | 0.066 | 0.211 | 42 | 0.315 | 0.754 |
|  | Distance of closest approach | -7.778*10<sup>-4</sup> | 0.003 | 42 | -0.243 | 0.809 |
|  | Duration of trial | 0.002 | 0.001 | 42 | 1.157 | 0.254 |
|  | **Soc. Env.** |  |  |  |  |  |
|  | **Cooperation** | **0** | **-** | **42** | **-** | **-** |
|  | **Defection** | **0.301** | **0.131** | **42** | **2.295** | **0.027** |
|  | Experimental Condition |  |  |  |  |  |
|  | Control (plastic plant) | 0 | - | 42 | - | - |
|  | Predator | 0.214 | 0.136 | 42 | 1.576 | 0.123 |
|  | **Soc. Env. x Exp. Condition** |  |  |  |  |  |
|  | **Cooperation - Control** | **0** | **-** | **42** | **-** | **-** |
|  | **Defection - Predator** | **-0.403** | **0.192** | **42** | **-2.095** | **0.042** |
| Mid-section | **Intercept** | **-2.016** | **0.490** | **42** | **-4.111** | **<0.001** |
|  | Standard body length | -0.008 | 0.205 | 42 | -0.039 | 0.969 |
|  | Distance of closest approach | -1.782*10<sup>-4</sup> | 0.003 | 42 | 0.063 | 0.950 |

| Brain section |  |  | Estimate | Standard error | df | z-value | p value |
| --- | --- | --- | --- | --- | --- | --- | --- |
|  | Duration of trial |  | 0.002 | 0.001 | 42 | 1.386 | 0.173 |
|  | Soc. Env. | Cooperation | 0 | - | 42 | - | - |
|  |  | Defection | 0.188 | 0.121 | 42 | 1.553 | 0.128 |
|  | Experimental Condition | Control (plastic plant) | 0 | - | 42 | - | - |
|  |  | Predator | 0.041 | 0.125 | 42 | 0.330 | 0.743 |
|  | Soc. Env. x Exp. Condition | Cooperation - Control | 0 | - | 42 | - | - |
|  |  | Defection - Predator | -0.257 | 0.179 | 42 | -1.438 | 0.158 |
| Hind-section | **Intercept** |  | **-2.455** | **0.512** | **40** | **-4.799** | **<0.001** |
|  | Standard body length |  | 0.084 | 0.207 | 40 | 0.407 | 0.687 |
|  | Distance of closest approach |  | -0.001 | 0.003 | 40 | -0.356 | 0.729 |
|  | Duration of trial |  | 6.218*10<sup>-5</sup> | 9.078*10<sup>-4</sup> | 40 | 0.068 | 0.946 |
|  | **Soc. Env.** | **Cooperation** | **0** | **-** | **40** | **-** | **-** |
|  |  | **Defection** | **0.341** | **0.124** | **40** | **2.662** | **0.011** |
|  | Experimental Condition | Control (plastic plant) | 0 | - | 40 | - | - |
|  |  | Predator | 0.145 | 0.124 | 40 | 1.175 | 0.247 |
|  | Soc. Env. x Exp. Condition | Cooperation - Control | 0 | - | 40 | - | - |

| Brain<br>section |  |  |  |  |  |  |
| --- | --- | --- | --- | --- | --- | --- |
|  |  | Estimate | Standard<br>error | df | z-value | p value |
|  | Defection - Predator | -0.286 | 0.187 | 40 | -1.535 | 0.133 |

**Table 6.** Post hoc analysis for the ‘Social Environment x Experimental Condition‘ interaction on the logarithm of the fore-section 5-HIAA/5-HT ratio. Pairwise least squares means comparisons after Tukey adjustment for multiple comparisons. Statistically significant contrasts are shown in bold.

| Contrast | Estimate | Standard error | df | t-value | p-value |
| --- | --- | --- | --- | --- | --- |
| Coop. Control – Def. Control | -0.301 | 0.131 | 42 | -2.295 | 0.116 |
| Coop. Control – Coop. Predator | -0.214 | 0.136 | 42 | -1.576 | 0.403 |
| Coop. Control – Def. Predator | -0.112 | 0.138 | 42 | -0.808 | 0.850 |
| Def. Control – Coop. Predator | 0.086 | 0.139 | 42 | 0.619 | 0.923 |
| Def. Control – Def. Predator | 0.189 | 0.137 | 42 | 1.376 | 0.521 |
| Coop. Predator – Def. Predator | 0.103 | 0.145 | 42 | 0.706 | 0.894 |

## 4 Discussion

Our results demonstrate that the cooperativeness of an individual’s social partners during predator inspection affects monoaminergic activity in the brain of female Trinidadian guppies. Dopaminergic and serotonergic neurotransmission were affected in the brain hind-section, as was brain fore-section serotonergic activity. Interestingly, we found no effect of ostensibly experiencing cooperation or defection in mid-section neurotransmission rates. To the best of our knowledge, this study provides the first insight into the role of brain monoaminergic neurotransmission systems in the experience of conspecific partner behaviour during cooperative interactions, and increases our understanding of the neural pathways potentially underlying conditional cooperative and affiliative behaviour among non-kin.

DA signalling has been implicated in reward and risk assessment [e.g. 37] and dopaminergic activity has been shown to play a role in the expression of cooperative behaviour in teleost fishes. For example, Messias and colleagues [35] found that disruption of dopamine neurotransmission in bluestreak cleaner wrasse (*Labroides dimidiatus*) resulted in increased cooperative effort, as shown by the increased frequency of costly behaviours usually linked to reconciliation after cheating client reef fish. We found that the effect of predator inspection on DOPAC/DA turnover rates was modified by the ostensible presence of a cooperative partner: when fish experienced defection from their social environment, predator inspection led to decreased hind-section DOPAC/DA turnover rates compared to our control condition (investigation of a familiar object). Larger groups of inspecting fish provide safety due to the dilution of risk [64], as well as their increased ability to detect and avoid predators [65, 66]; consequently, the presence of a cooperative partner during predator inspection is expected to decrease the risk of predation for each inspecting fish. It is therefore likely that fish who inspected a predator on their own (i.e. experimental defection) perceived the highest level of risk compared to the other experimental groups. Overall, the result supports the expectation that experiencing defection during predator inspection is likely to affect downstream behaviour in a way not induced by the experience of cooperation [67].

Hind-section HVA/DA ratios were affected by the ostensible behaviour of social partners irrespective of whether they were in a cooperative context or not (experimental predator stimulus or control familiar plant stimulus, respectively), with fish swimming in the lane as a singleton showing increased HVA/DA ratios compared to those ostensibly swimming in the lane pairwise. DA can be metabolised to either DOPAC after deamination by monoamine oxidase, or to 3-methoxy-tyramine (3-MT) after methylation by catechol-*O*-methyl transferase (COMT); both metabolites can be further converted to HVA, with the importance of each pathway being species-dependent [22]. The effects of social partner behaviour on hind-section HVA/DA ratio may thus be time-dependent and difficult to interpret independently of DOPAC/DA rates; more research into the importance of the DOPAC and 3-MT pathways in this species is needed to understand their involvement in behaviour.

We observed an effect of the interaction of experimental condition (predator versus no predator control) and the social environment (ostensible experience of cooperation versus defection) on fore-section 5-HIAA/5-HT ratios, suggesting that the serotonergic system is involved in the appraisal of partner behaviour during inspection. While post hoc analysis did not reveal any statistically significant differences between specific experimental conditions due to the lack of statistical power, our data suggest that the response to experiencing cooperation or defection differs depending on the context (presence of a predator). In guppies, phasic serotonin has been demonstrated to increase motivation to participate in predator inspection, while tonic serotonin increases cooperative behaviour in this paradigm [53], supporting the involvement of this system in the expression of behaviour in this cooperative context. Serotonin has also been shown to affect heterospecific cooperation between bluestreak cleaner wrasse and client reef fish by increasing the motivation and the probability of approaching a client fish, without affecting cleaning quality [40] – an effect probably mediated by an increased perception of risk posed by the client fish [34, 68]. The serotonergic system also has a well-documented role in stress responses [25, 69], with serotonergic activity increasing as a result of predator exposure [59, 70]. Given that shoaling acts as a mechanism of reducing risk of predation [64, 66], the differences in fore-section serotonergic activity may reflect differences in risk perception due to the presence or absence of a conspecific during forays away from the shoal.

The presence of conspecifics has been demonstrated to down-regulate responses to a detected threat – a phenomenon known as social buffering [71–74]. It is possible that social buffering occurs in inspection groups (and to a greater extent in larger groups), reducing the stress of approaching and inspecting a potential predator. Research points to the lateral amygdala (LA), the central amygdala (CeA) and the hypothalamic paraventricular nucleus (PVN) [75–78] as the neural substrate of social buffering in mammals; in teleosts, social buffering has been demonstrated to involve the medial part of the dorsal telencephalon (Dm – the teleostean homologue of the basolateral amygdala), the suppracommissural part of the ventral pallium (Vs – homologous to the extended amygdala) and the preoptic area (POA) [74]. As the Dm, Vs, and POA are located within the forebrain [79, 80], it is possible that the differences in fore-section serotonergic activity observed here reflect the effect of social buffering, where the presence of a conspecific (i.e. the experience of cooperation) induces a more positive affective state than a partner’s absence and that it is moderated to some extent by the level of threat (i.e. proximate predator cues present or absent). This finding is in accordance with the well documented role of the serotonergic system in stress [25, 69] and the increased risk of predation undertaken by lone inspectors [81].

Contrary to the fore-section, serotonergic activity in the hind-section was affected only by the ostensible behaviour of social partners, irrespective of whether there were proximate cues of a predator or not. A similar effect was observed in whole brain serotonergic activity, as fish swimming in the lane and approaching the stimulus compartment as a singleton showed increased 5-HIAA/5-HT ratios compared to those approaching as a pair, irrespective of whether this was a cooperative context or not. Hind-brain serotonergic activity has been linked to agonistic behaviour in teleosts [25, 26], and in particular the formation of dominance hierarchies [32, 82]; it is therefore likely that serotonin neurotransmission in this brain section mainly plays a role in the encoding of social stimuli across the manipulated social contexts.

While the effects observed here may be related to the effects of risk perception, our findings suggest that, at least in some brain sections, neurotransmission also encodes the cooperative behaviour of social partners in this context. The presence of a social partner affected neurotransmission differentially depending on the level of risk perceived. It is likely that the effect of the proximity of social partners on neurotransmission reflects social buffering and overall experiencing support from the social environment [83, 84]. Given the involvement of monoamine neurotransmission in reward [18, 19] and the characterisation of social stimuli [1], it is possible that the neurotransmission profiles elicited by experiences of cooperation or defection reflect the valence and salience of these social experiences. As monoamine neurotransmitters have been shown to affect the expression of cooperative behaviour, it is likely that these differences in internal state will mediate the appropriate behavioural responses to experiencing cooperation or defection [4], such as the decision to reciprocate the behaviour of social partners.

Overall, our data suggest that monoaminergic activity is largely affected by the cooperative behaviour of social partners (i.e. experiencing cooperation or defection); in some instances, such as serotonergic activity in the hind-section, this was irrespective of the inspection stimulus (i.e. the presence of a predator). Given our methodology, it is possible that these findings are not necessarily reflecting the effects of performing an inspection as a singleton or with a social partner on brain monoaminergic activity, but effects that arise from the use of the mirror itself. For example, it could be that the difference in the perceived environment (e.g. the lane size) between the two social conditions is affecting monoaminergic activity. Another difficulty in the interpretation of the difference between the experimental (predator) and control (plant) condition arises from the fact that the presence of a social partner would be perceived as cooperation in the former but not in the latter case, given that the aquarium plant is a familiar, not threatening, stimulus. In this sense, being joined by a simulated social partner in the absence of a predator (control condition) controls for the membership of social group. Our findings are supported by previous work by Pimentel and colleagues [53], who using the mirror paradigm also found evidence supporting the role of the serotonergic system in cooperative behaviour during predator inspection in guppies. They are also supported by recent work using a similar experimental setup, reporting an effect of experiencing cooperation or defection in brain oxytocin activity in a high predation population of Trinidadian guppies [85]. More work is needed to explicitly test the effects that the use of mirrors, as well as the perceived level of threat have on monoaminergic neurotransmission.

Here, we show that monoaminergic neurotransmission is affected by social experiences during predator inspection in female Trinidadian guppies. The activity of both the dopaminergic and serotonergic systems differed among brain sections. Given the involvement of these systems in a wide array of functions, such as prediction error, associative learning and social buffering, there are a number of possible drivers underlying the effects found in this study. Overall, however, the different neurotransmission patterns observed here are indicative of the effect of the experience of cooperative and non-cooperative social partners on an individual’s internal or affective state, and are thus likely to contribute to determining subsequent behavioural response to these experiences [4], and, ultimately, to the patterning of cooperative interactions among individuals in the population.

### 5 Conclusions

We found that in a laboratory population of female Trinidadian guppies originating from a high predation habitat, experiencing cooperation or defection from their social environment during predator inspection affected dopaminergic and serotonergic activity across the brain. Our findings provide further insight into the neuromodulation of underlying behavioural response to social experiences.

## Acknowledgments

The authors would like to thank Christine Soper, Adam Johnstone and Stephanie Fox for their help with animal husbandry.

## Declarations of interest

The authors declare no competing interests.

## Authors’ contributions

**Sylvia Dimitriadou:** Conceptualization, Methodology, Investigation, Formal analysis, Writing – Original Draft, Visualization; **Svante Winberg:** Resources, Methodology, Writing – Reviewing and Editing; **Per-Ove Thörnqvist:** Methodology, Investigation, Writing – Reviewing and Editing; **Darren Croft:** Conceptualization, Writing – Reviewing and Editing; **Safi Darden:** Conceptualization, Methodology, Investigation, Supervision, Funding acquisition, Writing – Reviewing and Editing

## Funding

This study was funded by the Danish Council for Independent Research (DFF – 1323-00105).

## Data availability statement

All data generated in this study are included in this manuscript; the code and data will be uploaded to Figshare upon acceptance.

## List of abbreviations used

DA: Dopamine
NE: Norepinephrine
5-HT: 5-hydroxytryptamine
5-HIAA: 5-hydroxyindoleacetic acid
DOPAC: 3,4-dihydroxyphenylacetic acid
HVA: Homovanillic acid
3-MT: 3-methoxy-tyramine
COMT: Catechol-O-methyl transferase
LA: Lateral amygdala
CeA: Central amygdala
PVN: Paraventricular nucleus
Dm: Dorsal telencephalon
Vs: Suppracommissural part of the ventral pallium
POA: Preoptic area

## Notes

### Competing Interest Statement

The authors have declared no competing interest.

## References

1. A.I. Faustino, G.A. Oliveira, R.F. Oliveira, Linking appraisal to behavioral flexibility in animals: implications for stress research, Front. Behav. Neurosci. 9 (2015) 104. https://doi.org/10.3389/fnbeh.2015.00104.

2. M. Kosfeld, M. Heinrichs, P.J. Zak, U. Fischbacher, E. Fehr, Oxytocin increases trust in humans, Nature. 435 (2005) 673–676. https://doi.org/10.1038/nature03701.

3. M. Cerqueira, S. Millot, M.F. Castanheira, A.S. Félix, T. Silva, G.A. Oliveira, C.C. Oliveira, C.I.M. Martins, R.F. Oliveira, Cognitive appraisal of environmental stimuli induces emotion-like states in fish, Sci. Rep. 7 (2017) 13181. https://doi.org/10.1038/s41598-017-13173-x.

4. M. Mendl, E.S. Paul, Animal affect and decision-making, Neurosci. Biobehav. Rev. 112 (2020) 144–163. https://doi.org/10.1016/j.neubiorev.2020.01.025.

5. A. Rangel, C. Camerer, P.R. Montague, A framework for studying the neurobiology of value-based decision making, Nat. Rev. Neurosci. 9 (2008) 545–556. https://doi.org/10.1038/nrn2357.

6. J.L. Sachs, U.G. Mueller, T.P. Wilcox, J.J. Bull, The evolution of cooperation, Q. Rev. Biol. 79 (2004) 135–160.

7. T. Clutton-Brock, Cooperation between non-kin in animal societies, Nature. 462 (2009) 51–57. https://doi.org/10.1038/nature08366.

8. L. Lehmann, L. Keller, The evolution of cooperation and altruism – a general framework and a classification of models, J. Evol. Biol. 19 (2006) 1365–1376. https://doi.org/10.1111/j.1420-9101.2006.01119.x.

9. S.A. West, A.S. Griffin, A. Gardner, Evolutionary Explanations for Cooperation, Curr. Biol. 17 (2007) 661–672. https://doi.org/10.1016/j.cub.2007.06.004.

10. M.A. Nowak, S. Roch, Upstream reciprocity and the evolution of gratitude, Proc. R. Soc. B Biol. Sci. 274 (2007) 605–610. https://doi.org/10.1098/rspb.2006.0125.

11. C.A. Aktipis, When to walk away and when to stay: cooperation evolves when agents can leave unproductive partners and groups, (2008).

12. C. Rutte, M. Taborsky, The influence of social experience on cooperative behaviour of rats (Rattus norvegicus): Direct vs generalised reciprocity, Behav. Ecol. Sociobiol. 62 (2008) 499–505. https://doi.org/10.1007/s00265-007-0474-3.

13. C. Rutte, M. Taborsky, Generalized reciprocity in rats, PLoS Biol. 5 (2007) e196.

14. L.A. Dugatkin, M. Alfieri, Guppies and the TIT FOR TAT strategy: preference based on past interaction, Behav. Ecol. Sociobiol. 28 (1991) 243–246. https://doi.org/10.1007/BF00175096.

15. S.K. Darden, R. James, J.M. Cave, J.B. Brask, D.P. Croft, Trinidadian guppies use a social heuristic that can support cooperation among non-kin, Proc. R. Soc. B Biol. Sci. 287 (2020) 20200487. https://doi.org/10.1098/rspb.2020.0487.

16. W. Schultz, Multiple Dopamine Functions at Different Time Courses, Annu. Rev. Neurosci. 30 (2007) 259–288. https://doi.org/10.1146/annurev.neuro.28.061604.135722.

17. B.P. Ramos, A.F.T. Arnsten, Adrenergic pharmacology and cognition: Focus on the prefrontal cortex, Pharmacol. Ther. 113 (2007) 523–536. https://doi.org/10.1016/J.PHARMTHERA.2006.11.006.

18. K.C. Berridge, T.E. Robinson, What is the role of dopamine in reward: Hedonic impact, reward learning, or incentive salience?, Brain Res. Rev. 28 (1998) 309–369. https://doi.org/10.1016/S0165-0173(98)00019-8.

19. J.D. Salamone, M. Correa, The mysterious motivational functions of mesolimbic dopamine, Neuron. 76 (2012) 470–485.

20. W. Schultz, Predictive reward signal of dopamine neurons., J. Neurophysiol. 80 (1998) 1–27. http://www.ncbi.nlm.nih.gov/pubmed/9658025 (accessed September 30, 2017).

21. F. Chaouloff, Serotonin, stress and corticoids, J. Psychopharmacol. 14 (2000) 139–151. https://doi.org/10.1177/026988110001400203.

22. C. Sørensen, I.B. Johansen, Ø. Øverli, Physiology of social stress in fishes, Physiol. Fishes (Eds DH Evans, JB Clairborne, S Currie). (2013) 289.

23. J.G. Hensler, Serotonin in Mood and Emotion, Handb. Behav. Neurosci. 21 (2010) 367–378. https://doi.org/10.1016/S1569-7339(10)70090-4.

24. D.E.A. Bush, E.M. Caparosa, A. Gekker, J. LeDoux, Beta-Adrenergic Receptors in the Lateral Nucleus of the Amygdala Contribute to the Acquisition but Not the Consolidation of Auditory Fear Conditioning, Front. Behav. Neurosci. 4 (2010) 154. https://doi.org/10.3389/fnbeh.2010.00154.

25. S. Winberg, G.E. Nilsson, Roles of brain monoamine neurotransmitters in agonistic behaviour and stress reactions, with particular reference to fish, Comp. Biochem. Physiol. Part C Comp. 106 (1993) 597–614. https://doi.org/10.1016/0742-8413(93)90216-8.

26. S. Winberg, P.O. Thörnqvist, Role of brain serotonin in modulating fish behavior, Curr. Zool. 62 (2016) 317–323. https://doi.org/10.1093/cz/zow037.

27. T. Canli, K.-P. Lesch, Long story short: the serotonin transporter in emotion regulation and social cognition, Nat. Neurosci. 10 (2007) 1103–1109. https://doi.org/10.1038/nn1964.

28. E. Höglund, F.A. Weltzien, J. Schjolden, S. Winberg, H. Ursin, K.B. Døving, Avoidance behavior and brain monoamines in fish, Brain Res. 1032 (2005) 104–110. https://doi.org/10.1016/j.brainres.2004.10.050.

29. C.H. Summers, S. Winberg, Interactions between the neural regulation of stress and aggression, J. Exp. Biol. 209 (2006) 4581LP – 4589. http://jeb.biologists.org/content/209/23/4581.abstract.

30. S. Winberg, Ø. Øverli, O. Lepage, Suppression of aggression in rainbow trout (Oncorhynchus mykiss) by dietary L-tryptophan, J. Exp. Biol. 204 (2001) 3867 LP – 3876. http://jeb.biologists.org/content/jexbio/204/22/3867.full.pdf (accessed September 30, 2017).

31. S.J. Dahlbom, T. Backström, K. Lundstedt-Enkel, S. Winberg, Aggression and monoamines: Effects of sex and social rank in zebrafish (Danio rerio), Behav. Brain Res. 228 (2012) 333–338. https://doi.org/10.1016/J.BBR.2011.12.011.

32. S. Winberg, G.E. Nilsson, K.H. Olsen, Social rank and brain levels of monoamines and monoamine metabolites in Arctic charr, Salvelinus alpinus (L.), J. Comp. Physiol. A. 168 (1991) 241–246. https://doi.org/10.1007/BF00218416.

33. P.J.J. Baarendse, C.A. Winstanley, L.J.M.J. Vanderschuren, Simultaneous blockade of dopamine and noradrenaline reuptake promotes disadvantageous decision making in a rat gambling task, Psychopharmacology (Berl). 225 (2013) 719–731. https://doi.org/10.1007/s00213-012-2857-z.

34. M.C. Soares, J.R. Paula, R. Bshary, Serotonin blockade delays learning performance in a cooperative fish, Anim. Cogn. 19 (2016) 1027–1030. https://doi.org/10.1007/s10071-016-0988-z.

35. J.P.M. Messias, J.R. Paula, A.S. Grutter, R. Bshary, M.C. Soares, Dopamine disruption increases negotiation for cooperative interactions in a fish, Sci. Rep. 6 (2016) 20817. https://doi.org/10.1038/srep20817.

36. J.D. Salamone, M. Correa, S.M. Mingote, S.M. Weber, A.M. Farrar, Nucleus Accumbens Dopamine and the Forebrain Circuitry Involved in Behavioral Activation and Effort-Related Decision Making: Implications for Understanding Anergia and Psychomotor Slowing in Depression, Curr. Psychiatry Rev. (2006). https://s3.amazonaws.com/academia.edu.documents/44264789/Nucleus_Accumbens_Dopamine_and_the_Foreb20160331-19792-ppprvy.pdf?AWSAccessKeyId=AKIAIWOWYYGZ2Y53UL3A&Expires=1522846149&Signature=EFatUOlHR%2B00kIaGCIGiq1JoQM4%3D&response-content-disposition=inlin (accessed April 4, 2018).

37. W. Schultz, Dopamine signals for reward value and risk: basic and recent data, Behav. Brain Funct. 6 (2010) 24. https://doi.org/10.1186/1744-9081-6-24.

38. P.W. Glimcher, Understanding dopamine and reinforcement learning: the dopamine reward prediction error hypothesis., Proc. Natl. Acad. Sci. U. S. A. 108 Suppl 3 (2011) 15647–54. https://doi.org/10.1073/pnas.1014269108.

39. J. LeDoux, Rethinking the Emotional Brain, Neuron. 73 (2012) 653–676. https://doi.org/10.1016/J.NEURON.2012.02.004.

40. J.R. Paula, J.P. Messias, A.S. Grutter, R. Bshary, M.C. Soares, The role of serotonin in the modulation of cooperative behavior, Behav. Ecol. 26 (2015) 1005–1012. https://doi.org/10.1093/beheco/arv039.

41. M.C. Soares, S.C. Cardoso, J.T. Malato, J.P.M. Messias, Can cleanerfish overcome temptation? A selective role for dopamine influence on cooperative-based decision making, Physiol. Behav. 169 (2017) 124–129. https://doi.org/10.1016/j.physbeh.2016.11.028.

42. T.J. Pitcher, D.A. Green, A.E. Magurran, Dicing with death: predator inspection behaviour in minnow shoals, J. Fish Biol. 28 (1986) 439–448. https://doi.org/10.1111/j.1095-8649.1986.tb05181.x.

43. J.R. Allan, T.J. Pitcher, Species segregation during predator evasion in cyprinid fish shoals, Freshw. Biol. 16 (1986) 653–659.

44. A.E. Magurran, B.H. Seghers, Predator inspection behaviour covaries with schooling tendency amongst wild guppy, Poecilia reticulata, populations in Trinidad, Behaviour. 128 (1994) 121–134.

45. M. Milinski, Tit for tat in sticklebacks and the evolution of cooperation, Nature. 325 (1987) 433–435.

46. S. Dimitriadou, E.M. Santos, D.P. Croft, R. van Aerle, I.W. Ramnarine, A.L. Filby, S.K. Darden, Social partner cooperativeness influences brain oxytocin transcription in Trinidadian guppies (Poecilia reticulata), Behav. Brain Res. (2021) 113643. https://doi.org/10.1016/J.BBR.2021.113643.

47. M. Edenbrow, B.H. Bleakley, S.K. Darden, C.R. Tyler, I.W. Ramnarine, D.P. Croft, The Evolution of Cooperation: Interacting Phenotypes among Social Partners, Am. Nat. 189 (2017) 000–000. https://doi.org/10.1086/691386.

48. L.A. Dugatkin, Tendency to inspect predators predicts mortality risk in the guppy (Poecilia reticulata), 3 (1992) 124–127.

49. A. De Santi, V.A. Sovrano, A. Bisazza, G. Vallortigara, Mosquitofish display differential left-and right-eye use during mirror image scrutiny and predator inspection responses, Anim. Behav. 61 (2001) 305–310.

50. L.A. Dugatkin, Do guppies play TIT FOR TAT during predator inspection visits?, Behav. Ecol. Sociobiol. 23 (1988) 395–399.

51. J.B. Brask, D.P. Croft, M. Edenbrow, R. James, B.H. Bleakley, I.W. Ramnarine, R.J.P. Heathcote, C.R. Tyler, P.B. Hamilton, T. Dabelsteen, S.K. Darden, Evolution of non-kin cooperation: social assortment by cooperative phenotype in guppies, R. Soc. Open Sci. 6 (2019) 181493. https://doi.org/10.1098/rsos.181493.

52. L.A. Dugatkin, M. Alfieri, Tit-For-Tat in guppies (Poecilia reticulata): the relative nature of cooperation and defection during predator inspection, Evol. Ecol. 5 (1991) 300– 309. https://doi.org/10.1007/BF02214234.

53. A.F.N. Pimentel, T. dos S. Carvalho, F. Lima, M. Lima-Maximino, M.C. Soares, C. Maximino, Conditional approach as cooperation in predator inspection: A role for serotonin?, Behav. Brain Res. 365 (2019) 164–169. https://doi.org/10.1016/J.BBR.2019.03.005.

54. B.H. Seghers, Analysis of geographic variation in the antipredator adaptations of the guppy : Poecilia reticulata, (1973). https://doi.org/10.14288/1.0100947.

55. A.E. Magurran, Evolutionary ecology: the Trinidadian guppy, Oxford University Press on Demand, 2005. https://doi.org/10.1093/acprof.

56. A.E. Magurran, S.L. Girling, Predator model recognition and response habituation in shoaling minnows, Anim. Behav. 34 (1986) 510–518.

57. L.A. Dugatkin, J.-G.J. Godin, Prey approaching predators: a cost-benefit perspective, in: Ann. Zool. Fennici, JSTOR, 1992: pp. 233–252. https://doi.org/10.2307/23735625.

58. P.O. Thörnqvist, E. Höglund, S. Winberg, Natural selection constrains personality and brain gene expression differences in Atlantic salmon (Salmo salar), J. Exp. Biol. 218 (2015) 1077 LP – 1083. http://jeb.biologists.org/content/218/7/1077.abstract.

59. A.M. Bell, T. Backström, F.A. Huntingford, T.G. Pottinger, S. Winberg, Variable neuroendocrine responses to ecologically-relevant challenges in sticklebacks, Physiol. Behav. 91 (2007) 15–25. https://doi.org/10.1016/j.physbeh.2007.01.012.

60. N.J. Shannon, J.W. Gunnet, K.E. Moore, A comparison of biochemical indices of 5-hydroxytryptaminergic neuronal activity following electrical stimulation of the dorsal raphe nucleus, J. Neurochem. 47 (1986) 958–965.

61. F. Cribari-Neto, A. Zeileis, Beta Regression in *R*, J. Stat. Softw. 34 (2010) 1–24. https://doi.org/10.18637/jss.v034.i02.

62. S. Ferrari, F. Cribari-Neto, Beta Regression for modelling rates and proportions, J. Appl. Stat. 31 (2004). https://pdfs.semanticscholar.org/09b8/ab5c04d6a914ae9a7157f7de8880205165fd.pdf (accessed November 17, 2017).

63. R.C. Team, R: A language and environment for statistical computing. Vienna, Austria: R Foundation for Statistical Computing; 2014, (2014).

64. T.J. Pitcher, Functions of Shoaling Behaviour in Teleosts, in: Behav. Teleost Fishes, Springer US, Boston, MA, 1986: pp. 294–337. https://doi.org/10.1007/978-1-4684-8261-4_12.

65. G. Roberts, Why individual vigilance declines as group size increases, Anim. Behav. 51 (1996) 1077–1086.

66. J. Krause, G.D. Ruxton, Living in groups, Oxford University Press, 2002.

67. M. Mendl, O.H.P. Burman, E.S. Paul, An integrative and functional framework for the study of animal emotion and mood., Proceedings. Biol. Sci. 277 (2010) 2895–904. https://doi.org/10.1098/rspb.2010.0303.

68. M.C. Soares, The Neurobiology of Mutualistic Behavior: The Cleanerfish Swims into the Spotlight, Front. Behav. Neurosci. 11 (2017) 191. https://doi.org/10.3389/fnbeh.2017.00191.

69. J.I. Johnsson, S. Winberg, K.A. Sloman, Social interactions, Fish Physiol. 24 (2006) 151.

70. S. Winberg, A.A.J. Myrberg, G.E. Nilsson, Predator exposure alters brain serotonin metabolism in bicolour damselfish., Neuroreport. 4 (1993) 399–402.

71. A.S. Smith, Z. Wang, Hypothalamic oxytocin mediates social buffering of the stress response., Biol. Psychiatry. 76 (2014) 281–8. https://doi.org/10.1016/j.biopsych.2013.09.017.

72. J. Edgar, S. Held, E. Paul, I. Pettersson, R. I’Anson Price, C. Nicol, Social buffering in a bird, Anim. Behav. 105 (2015) 11–19. https://doi.org/10.1016/J.ANBEHAV.2015.04.007.

73. M.B. Hennessy, S. Kaiser, N. Sachser, Social buffering of the stress response: Diversity, mechanisms, and functions, Front. Neuroendocrinol. 30 (2009) 470–482. https://doi.org/10.1016/J.YFRNE.2009.06.001.

74. A.I. Faustino, A. Tacão-Monteiro, R.F. Oliveira, Mechanisms of social buffering of fear in zebrafish, Sci. Rep. 7 (2017) 44329. https://doi.org/10.1038/srep44329.

75. Y. Kiyokawa, A. Honda, Y. Takeuchi, Y. Mori, A familiar conspecific is more effective than an unfamiliar conspecific for social buffering of conditioned fear responses in male rats, Behav. Brain Res. 267 (2014) 189–193. https://doi.org/10.1016/J.BBR.2014.03.043.

76. F. Fuzzo, J. Matsumoto, Y. Kiyokawa, Y. Takeuchi, T. Ono, H. Nishijo, Social buffering suppresses fear-associated activation of the lateral amygdala in male rats: behavioral and neurophysiological evidence, Front. Neurosci. 9 (2015) 99. https://doi.org/10.3389/fnins.2015.00099.

77. Y. Takahashi, Y. Kiyokawa, Y. Kodama, S. Arata, Y. Takeuchi, Y. Mori, Olfactory signals mediate social buffering of conditioned fear responses in male rats, Behav. Brain Res. 240 (2013) 46–51. https://doi.org/10.1016/J.BBR.2012.11.017.

78. A.P. da Costa, A.E. Leigh, M.-S. Man, K.M. Kendrick, Face pictures reduce behavioural, autonomic, endocrine and neural indices of stress and fear in sheep., Proceedings. Biol. Sci. 271 (2004) 2077–84. https://doi.org/10.1098/rspb.2004.2831.

79. R. Bshary, S. Gingins, A.L. Vail, Social cognition in fishes, Trends Cogn. Sci. 18 (2014) 465–471. https://doi.org/10.1016/j.tics.2014.04.005.

80. E.K. Fischer, S.E. Westrick, L. Hartsough, K.L. Hoke, Differences In Neural Activity, But Not Behavior, Across Social Contexts In Guppies, Poecilia reticulata, BioRxiv. (2018) 265736. https://doi.org/10.1101/265736.

81. M. Milinski, J.H. Lüthi, R. Eggler, G.A. Parker, Cooperation under predation risk: experiments on costs and benefits, Proc. R. Soc. London B Biol. Sci. 264 (1997) 831– 837. https://doi.org/10.1098/rspb.1997.0116.

82. S. Winberg, G.E. Nilsson, K.H. Olsen, Changes in brain serotonergic activity during hierarchic behavior in Arctic charr (Salvelinus alpinus L.) are socially induced, J. Comp. Physiol. A. 170 (1992) 93–99. https://doi.org/10.1007/BF00190404.

83. F.S. Chen, R. Kumsta, B. von Dawans, M. Monakhov, R.P. Ebstein, M. Heinrichs, Common oxytocin receptor gene (OXTR) polymorphism and social support interact to reduce stress in humans., Proc. Natl. Acad. Sci. U. S. A. 108 (2011) 19937–42. https://doi.org/10.1073/pnas.1113079108.

84. A. Meyer-Lindenberg, G. Domes, P. Kirsch, M. Heinrichs, Oxytocin and vasopressin in the human brain: Social neuropeptides for translational medicine, Nat. Rev. Neurosci. 12 (2011) 524–538. https://doi.org/10.1038/nrn3044.

85. S. Dimitriadou, E. Santos, D.P. Croft, R. van Aerle, I.W. Ramnarine, A.L. Filby, S.K. Darden, Social partner cooperativeness influences brain oxytocin transcription in Trinidadian guppies (Poecilia reticulata), BioRxiv. (2021) 2021.03.01.433346. https://doi.org/10.1101/2021.03.01.433346.

