## Supplementary information for "Brain monoaminergic activity during predator inspection in the Trinidadian guppy (*Poecilia reticulata*)"

**Supplementary figures**

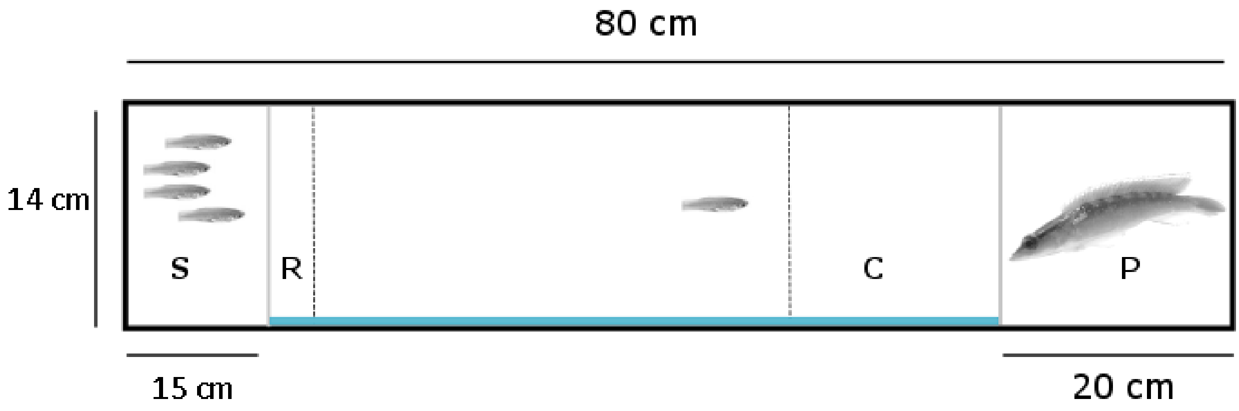


Fig. S1. The experimental setup used for the behavioural assay (top view). S: stimulus shoal compartment, perforated to allow for olfactory cue transmission. R: refuge area. C: area of close proximity to the predator. P: inspection stimulus compartment. A focal individual was placed in the inspection lane. A mirror was placed lengthwise to simulate cooperation (blue line); defection was simulated by an opaque partition.

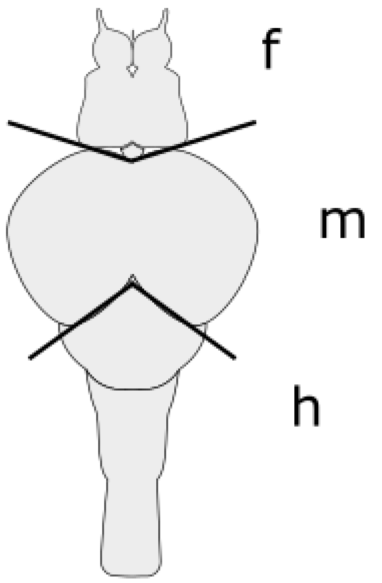


Fig. S2. Dorsal view of the brain of a female guppy. The black lines denote the brain section borders. f: fore-section; m: mid-section; h: hind-section. The brain outline is adapted from the SciDraw scientific figure preparation system (https://scidraw.io/)

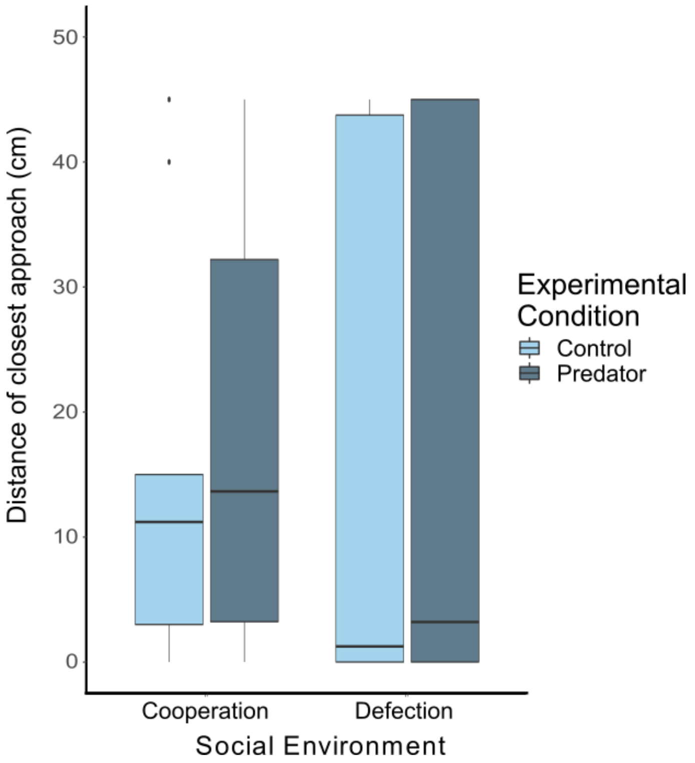


Fig. S3. Distance of closest approach (cm) to the stimulus compartment (Cooperation-Control: N= 10; Cooperation-Predator: N= 9; Defection-Control: N= 11; Defection-Predator: N= 8). Boxes represent the interquartile range (25th and 75th quartiles), and the horizontal lines represent the medians. The whiskers extend to the largest value (upper whisker) and lowest (lower whisker) value no further than 1.5 times the interquartile range. The dots represent outlying values.

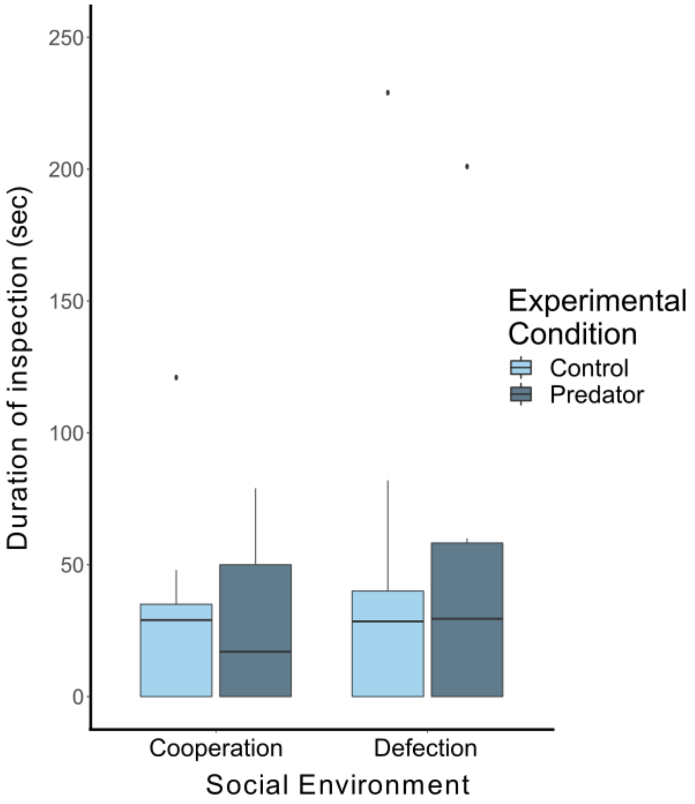


Fig. S4. Duration of the time (sec) spent inspecting the stimulus. (Cooperation-Control: N= 10; Cooperation-Predator: N= 9; Defection-Control: N= 11; Defection-Predator: N= 8). Boxes represent the interquartile range (25th and 75th quartiles), and the horizontal lines represent the medians. The whiskers extend to the largest value (upper whisker) and lowest (lower whisker) value no further than 1.5 times the interquartile range. The dots represent outlying values.
